# Sequestration and efflux largely account for cadmium and copper resistance in the deep sea epsilonproteobacterium, *Nitratiruptor* sp. SB155-2

**DOI:** 10.1101/2021.09.06.459102

**Authors:** Ángela Ares, Sanae Sakai, Toshio Sasaki, Satoshi Mitarai, Takuro Nunoura

## Abstract

In deep sea hydrothermal vent environments, metal- and metalloid-enriched fluids and sediments abound, making these habitats ideal to study metal resistance in prokaryotes. In this investigation, the architecture of the epsilonproteobacterium, *Nitratiruptor* sp. SB155-2 transcriptome in combination with sub-cellular analysis using scanning transmission electron microscopy and energy-dispersive X-ray spectroscopy (STEM-EDX) was examined to better understand mechanisms of tolerance for cadmium (Cd) and copper (Cu) at stress-inducing concentrations. Transcriptomic expression profiles were remarkably different in the presence of these two metals, displaying 385 (19%) and 629 (31%) genes differentially expressed (DE) in the presence of Cd and Cu, respectively, while only 7% of DE genes were shared, with genes for non-specific metal transporters and genes involved in oxidative stress-response predominating. The principal metal-specific DE pathways under Cu stress, including those involving sulfur, cysteine, and methionine, are likely required for high-affinity efflux systems, while flagella formation and chemotaxis were over-represented under Cd stress. Consistent with these differences, STEM-EDX analysis revealed that polyphosphate-like granules (pPLG), the formation of CdS particles, and the periplasmic space may be crucial for Cd sequestration. Overall, this study provides new insights regarding metal-specific adaptations of Epsilonproteobacteria to deep sea hydrothermal vent environments.

**Significance originality statement:** Deep sea hydrothermal vents are unique environments in which metals and metalloids abound. Despite being a dominant phylum in these environments, adaptations enabling Epsilonproteobacteria to thrive in metal-rich environments remain poorly understood. In this study, a combination of high-throughput, whole-transcriptome RNA-seq analysis, scanning transmission electron microscopy, and energy-dispersive X-ray spectroscopy provide a comprehensive picture of molecular and morphological adaptations controlling metal efflux and sequestration systems of this bacterium in response to cadmium and copper. Many of these responses are metal-specific.

## 1. Introduction

Bacteria are widely recognized for efficient adaptation to environmental stress, including adaptation to high concentrations of metals and metalloids. Unlike eukaryotes, bacteria lack compartments to store metal ions; hence, cellular homeostasis relies on metal ion uptake, efflux, and sequestration (Nies, 2003; Ma *et al*., 2009; Chandrangsu *et al*., 2017) to regulate the net accumulation of metals in the cytoplasm. Metals such copper, manganese, nickel or zinc are required for biological processes in only trace quantities (Reyes-Caballero *et al*., 2011), while others like silver, cadmium, and lead have no known biological function, so that it is crucial to remove them from the cytosol immediately. Metal ion uptake and efflux are commonly regulated by metalloregulatory proteins and high-affinity transporters that respond to various metal ions in highly selective ways. Predominant among efflux systems responsible for copper (Cu) homeostasis are the resistance-nodulation-division (RND) protein superfamily *cus* determinant (Franke *et al*., 2003) and P-type ATPase CopA, which pumps Cu ions out of the cell (Outten *et al*., 2000; Fan *et al*., 2001) in combination with multicopper oxidase CueO (Grass and Rensing, 2001). In contrast, under cadmium (Cd) stress, different systems operate, such as the genetic determinants, *cad* and *czc*, which export excess Cd, as well as other ions, like zinc or lead (Nucifora *et al*., 1989; Nies, 1995; Naz *et al*., 2005; Ducret *et al*., 2020). Additionally, sequestration of metals inside or outside the cell constitutes another major strategy. For this purpose, polyphosphate-like granules (pPLGs), long-chain anionic polymers of phosphate, constitute a non-selective strategy employed by many bacteria species (Keasling and Hupf, 1996; Villagrasa *et al*., 2020). Formation and hydrolysis of pPLGs are mainly due to the action of polyphosphate kinases (PPKs) and exopolyphosphatases (PPXs), respectively, and their activities can be modulated by metal concentrations in the cytosol, among other factors (Docampo, 2006). When metal ions are present in the environment, pPLGs can efficiently sequester them, but in some species and with some metals, accumulation of metal ions also tends to stimulate PPX activity, promoting hydrolysis into inorganic phosphate molecules, so that metal–phosphate complexes can be transported out of the cells (Alvarez and Jerez, 2004; Rivero *et al*., 2018). Another means of reducing metal presence in the cytosol is through biomineralization of metals. A number of bacteria are able to precipitate metal-laden particles in the periplasmic space. Biomineralization has been observed when bacteria are exposed to a variety of metals. Cd is converted to cadmium sulfide particles (CdS) (Yang *et al*., 2015; Wang *et al*., 2017; Ma *et al*., 2020; Ma and Sun, 2021), lead is excreted as galena (PbS) or lead (II) phosphate (Pb_3_(PO_4_)_2_) (Zhang *et al*., 2019), and Cu is converted to chalcocite (Cu_2_S) (Kimber *et al*., 2020). Many strategies to mitigate metal stress are metal-specific; hence, performing comparative metal studies (Nies, 1995; Hu *et al*., 2005; Lu *et al*., 2017; Jiang *et al*., 2020) in the same experimental set-up, enables metal-specific mechanisms to be distinguished from generalized molecular pathways. Furthermore, under natural conditions, various metals co-occur in the same micro-habitat, as in deep-sea hydrothermal vents.

In deep-sea hydrothermal vent environments, metals and reduced gas-rich hydrothermal fluids form steep physico-chemical environments, and life must adapt such extreme environments (Reysenbach *et al*., 2000; Takai and Nakamura, 2010). However, knowledge about mechanisms of metal resistance in microbes associated with hydrothermal vent environments are limited, except for the case of the well-known hyperthermophilic archaeon genus, *Thermococcus* (Fukui *et al*., 2005; Lagorce *et al*., 2012).

Epsilonproteobacteria are some of the representative primary producers in hydrothermal ecosystems, and in fact, they account for 66-98% of the microorganisms associated with hydrothermal vent substrates (Lopez-Garcia *et al*., 2003; Takai *et al*., 2003; Vetriani *et al*., 2014). However, to date, adaptations enabling Epsilonproteobacteria to thrive in metal-rich environments remain poorly understood. In this study, we characterized transcriptomic responses and morphological adaptations controlling metal efflux and sequestration systems responding to Cd and Cu stress in the deep-sea hydrothermal vent bacterium, *Nitratiruptor* sp. SB155-2. *Nitratiruptor* sp. SB155-2, a gram-negative, chemolithoautotrophic epsilonproteobacterium isolated from the Okinawa Trough, is capable of growing under microaerobic and anaerobic conditions (Nakagawa et al., 2007). Genomic features of *Nitratiruptor* sp. SB155-2 strain revealed ≥20 responsive genes in a wide variety of metal transport systems, as well as detoxification mechanisms of heavy metals such as arsenic, cadmium, copper, manganese, or zinc (Nakagawa et al., 2007), but these were not studied in detail. Special attention was paid to distinguishing metal-specific vs. generalized strategies to mitigate metal stress. To achieve this, this study employed high-throughput scanning transmission electron microscopy (STEM) coupled with energy-dispersive X-ray spectroscopy (EDX) and whole-transcriptome RNA-seq analysis.

## 2. Results

### 2.1. Effect of sublethal metal concentrations on growth of *Nitratiruptor* sp. SB155-2

Growth of *Nitratiruptor* sp. SB155-2 was monitored for five days by measuring cell densities every 12 h in the presence of cadmium (Cd) (0.05, 0.1, and 0.5 mM), or copper (Cu) (0.01, 0.05, 0.1mM), compared with controls (Figure 1A). Cd and Cu were selected because both are found in Okinawa Trough deposits (Chen *et al*., 2000) and are well known for their high toxicity for various microbial groups, which make them model elements for microbial toxicology studies (Nies, 2003; Hu *et al*., 2005; Arguello *et al*., 2013). Additionally, these elements have different biological roles. Whereas Cu is an essential element, required as a redox cofactor in the catalytic centers of enzymes (Gort *et al*., 1999; Cobine *et al*., 2006), Cd serves no known function in living organisms.

**Figure 1.**
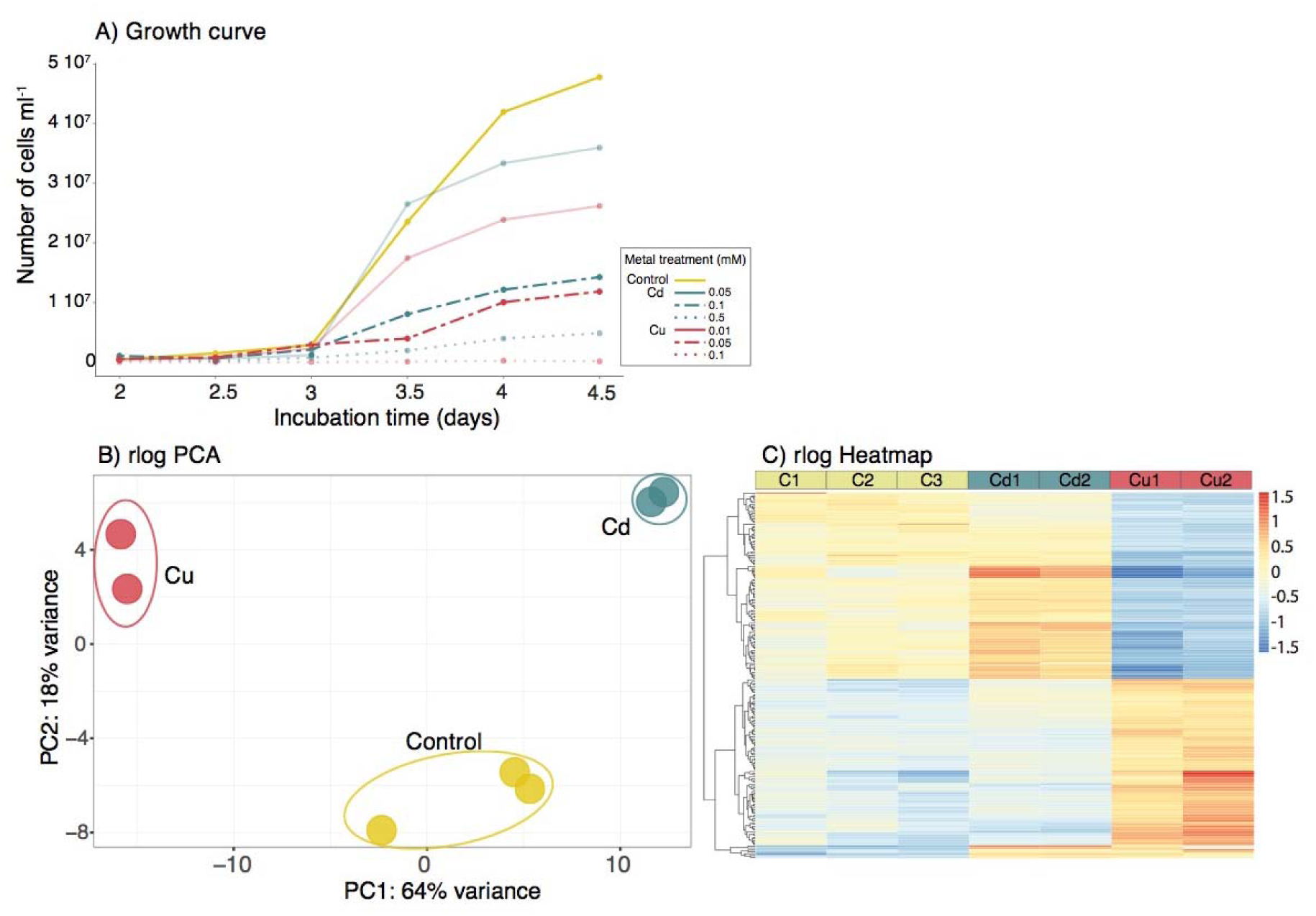
Growth performance of *Nitratiruptor* sp. SB155-2 cells affected by different metal concentrations and PCA and heatmap of RNA-seq transcriptome data demonstrated metal-specific gene expression patterns. **A)** Aliquots of 0.1 mL were collected from each bottle every 12 h for five days, stained with SYBR Green I and quantified using an Accuri C6 flow cytometer. Values are means of four replicates. Sub-lethal concentrations at which subsequent RNA-seq analysis and microscopy observations were performed are indicated with 100% opacity in the corresponding Control, Cd 0.1 mM, and Cu 0.05 mM. B) A PCA plot of all RNA-seq samples from regularized log transformed counts (rlog) showing how samples subjected to different treatments clustered separately. C) A heatmap including 300 genes with the lowest False Discovery Rate (FDR) adjusted p-values. Cell color represents differences from the mean regularized log transformed count for each contig in each sample. C= Controls; Cd= CdCl_2_ 0.1mM; Cu = CuSO_4_ 0.05 mM

In order to select meaningful stress conditions, the objective was to find the highest metal concentration that did not inhibit cell growth. For both elements, the medium concentration tested (Cd 0.1 mM and Cu 0.05 mM) impacted but did not inhibit the cell growth completely (Figure 1A), while allowing cell densities sufficient to perform RNAseq and TEM-EDX analysis.

### 2.2. Overview of RNA seq results

Illumina NovaSeqTM 6000 sequencing produced over 37.8 million read pairs with the number of read pairs per sample ranging from 4.9-6.6 million. After trimming, 37.4 million read pairs (∼ 99%) remained. Transcriptomic coverage of the *Nitratiruptor* sp. SB155-2 genome varied between 77.9-94% (Supplemental Table 1).

To determine whether various gene transcription profiles reflected different treatments, principal component analysis (PCA) was performed (Figure 1B). Sample variability was higher between different experimental treatments than between biological replicates and important differences were found between samples treated with Cd and Cu, with replicates clustering well separated along the main axis (64% of total variance). Overall expression patterns were evaluated using an expression heatmap (Figure 1C), which showed the most significant DE genes ordered by FDR-adjusted P-value (padj) for the first 200 genes. The resulting heatmap clearly illustrates different expression profiles in Cu-treated samples, compared with controls and Cd-treated samples.

### 2.3. Differentially expressed genes

Overall, the DESeq2 test identified 385 differentially expressed genes (DEGs) (19.3%) under Cd stress —with 190 and 195 genes significantly up- and down-regulated, respectively (Figure 2A). A more distinct response was found in samples treated with Cu, resulting in 31.6% of all total genes differentially expressed (i.e., 629 genes), 291 up-regulated and 338 down-regulated (Figure 2B). Accordingly, DEGs encoding transporter systems as well as oxidative stress responsive genes were found in higher numbers following Cu than Cd stress (Supplemental Table 3-4). For example, several genes encoding different efflux RND transporter subunits, ABC transporter ATP-binding protein, oxidoreductases, or thioredoxins/glutaredoxins were specifically up-regulated following Cd or Cu exposure.

**Figure 2.**
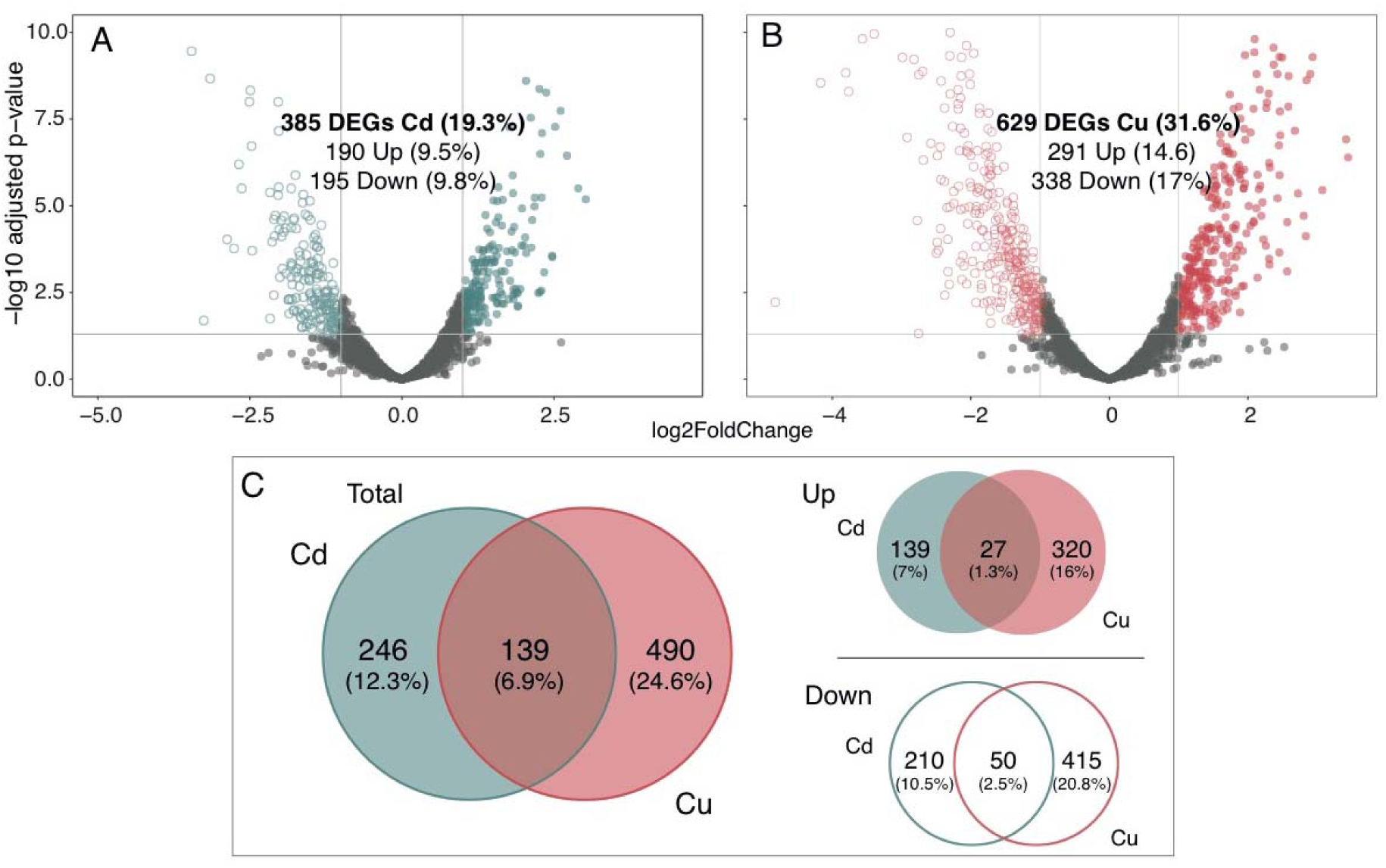
Metal-specific gene expression pattern in *Nitratiruptor* sp. SB155-2 with copper induced a significantly higher number of differentially expressed genes (DEGs) A-B) Volcano plots displaying gene expression patterns of *Nitratiruptor* sp. SB155-2 cells treated with cadmium (A) and copper (B) relative to controls. Significantly differentially expressed genes (DEGs) are highlighted in blue for Cd and red for Cu. C) Venn diagram showing DEGs occurring in both treatments. Log2FoldChange >1 or <-1, padj ≤ 0.05. C= Controls; Cd= CdCl_2_ 0.1mM; Cu = CuSO_4_ 0.05 mM.

Importantly, of the 139 DE genes that were expressed in the presence of both metals, only 77 were expressed with the same pattern (27 up- and 50 down-regulated), further demonstrating important differences in the *Nitratiruptor* sp. SB155-2 metal stress response resulting from exposure to Cd and Cu (Figure 2C). Table 1 shows up-regulated genes that responded to both Cd and Cu stress. Two groups of contiguous genes were clustered at two different chromosome locations, likely corresponding to two multicomponent systems to transport both Cd and Cu ions from the cytoplasm to the extracellular environment. The first group (i.e., NIS_RS00145, NIS_RS00150, NIS_RS00155), is formed by a metalloregulator ArsR/SmtB family transcription factor, a permease, and an arsenate reductase, ArsC, likely encoding the *ars* operon (Ben Fekih *et al*., 2018). Other genes involved in the *ars* operon, ArsB and ArsR, were upregulated following Cu and Cd exposure, respectively. The second group comprised five upregulated genes induced under Cd and Cu stress (i.e., NIS_RS04910, NIS_RS04915, NIS_RS04920, NIS_RS04930, NIS_RS04935) and six under Cd stress alone, with NIS_RS04925 likely encoding a multidrug efflux RND transporter. The genes forming this second system encode a SO_0444 family Cu/Zn efflux transporter, cytochrome c, a metalloregulator ArsR/SMtB family transcription factor, a hypothetical protein, and an outer membrane protein, TolC. Other operons commonly up-regulated included parts of the oxidative stress response, such as the GroES/GroEL or DnaK/DnaJ molecular chaperone systems.

**Table 1.**
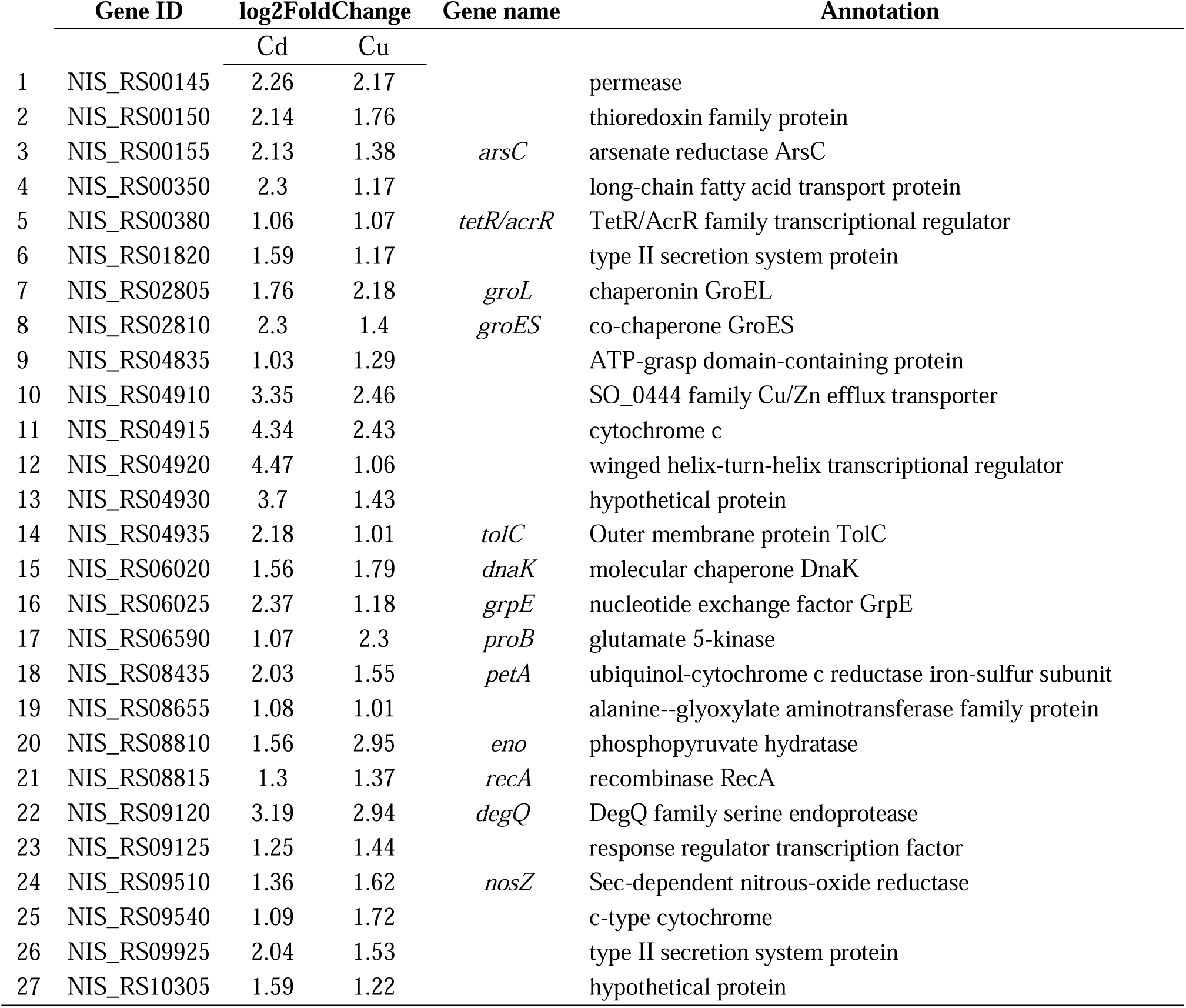
Differentially up-regulated genes of *Nitratiruptor sp*. SB155-2 respond to both cadmium (0.1 mM) and copper (0.05 mM) stress, showing the stress response and expression ratios (in log2FoldChange).

|  | Gene ID | log2FoldChange |  | Gene name | Annotation |
| --- | --- | --- | --- | --- | --- |
|  |  | Cd | Cu |  |  |
| 1 | NIS_RS00145 | 2.26 | 2.17 |  | permease |
| 2 | NIS_RS00150 | 2.14 | 1.76 |  | thioredoxin family protein |
| 3 | NIS_RS00155 | 2.13 | 1.38 | *arsC* | arsenate reductase ArsC |
| 4 | NIS_RS00350 | 2.3 | 1.17 |  | long-chain fatty acid transport protein |
| 5 | NIS_RS00380 | 1.06 | 1.07 | *tetR/acrR* | TetR/AcrR family transcriptional regulator |
| 6 | NIS_RS01820 | 1.59 | 1.17 |  | type II secretion system protein |
| 7 | NIS_RS02805 | 1.76 | 2.18 | *groL* | chaperonin GroEL |
| 8 | NIS_RS02810 | 2.3 | 1.4 | *groES* | co-chaperone GroES |
| 9 | NIS_RS04835 | 1.03 | 1.29 |  | ATP-grasp domain-containing protein |
| 10 | NIS_RS04910 | 3.35 | 2.46 |  | SO_0444 family Cu/Zn efflux transporter |
| 11 | NIS_RS04915 | 4.34 | 2.43 |  | cytochrome c |
| 12 | NIS_RS04920 | 4.47 | 1.06 |  | winged helix-turn-helix transcriptional regulator |
| 13 | NIS_RS04930 | 3.7 | 1.43 |  | hypothetical protein |
| 14 | NIS_RS04935 | 2.18 | 1.01 | *tolC* | Outer membrane protein TolC |
| 15 | NIS_RS06020 | 1.56 | 1.79 | *dnaK* | molecular chaperone DnaK |
| 16 | NIS_RS06025 | 2.37 | 1.18 | *grpE* | nucleotide exchange factor GrpE |
| 17 | NIS_RS06590 | 1.07 | 2.3 | *proB* | glutamate 5-kinase |
| 18 | NIS_RS08435 | 2.03 | 1.55 | *petA* | ubiquinol-cytochrome c reductase iron-sulfur subunit |
| 19 | NIS_RS08655 | 1.08 | 1.01 |  | alanine--glyoxylate aminotransferase family protein |
| 20 | NIS_RS08810 | 1.56 | 2.95 | *eno* | phosphopyruvate hydratase |
| 21 | NIS_RS08815 | 1.3 | 1.37 | *recA* | recombinase RecA |
| 22 | NIS_RS09120 | 3.19 | 2.94 | *degQ* | DegQ family serine endoprotease |
| 23 | NIS_RS09125 | 1.25 | 1.44 |  | response regulator transcription factor |
| 24 | NIS_RS09510 | 1.36 | 1.62 | *nosZ* | Sec-dependent nitrous-oxide reductase |
| 25 | NIS_RS09540 | 1.09 | 1.72 |  | c-type cytochrome |
| 26 | NIS_RS09925 | 2.04 | 1.53 |  | type II secretion system protein |
| 27 | NIS_RS10305 | 1.59 | 1.22 |  | hypothetical protein |

### 2.4. Enrichment analysis: Gene Ontology and KEGG pathways

The bioinformatic platform, Blast2GO, was used to annotate Gene Ontology terms associated with DEGs found in this study. From these, in order to identify over-represented GO terms involved in the *Molecular Function* (MF) and *Biological Process* (BP) categories under different metal treatments, the R package GOstats, was used (padj<0.05). Overall, 25 BP GO terms, 9 and 16, in the presence of Cd and Cu, respectively, and 29 MF GO terms, 19 and 10, in the presence of Cd and Cu, respectively, were over-represented. BP and MF GO terms were displayed using the REVIGO online tool for effective visualization in a multidimensional scaling plot ordered by semantic similarity (Supek et al. 2011) (Figure 3). Following Cu exposure, GO terms involving *ion channel, ion binding, sulfur compound biosynthesis, thioester metabolic process*, and *O-acetylhomoserine aminocarboxypropyltransferase* activity terms were over-represented. By contrast, following Cd exposure, other GO terms were over-represented, including *microtubule cytoskeleton organization, spindle assembly, gene expression, phosphorelay signal transduction system, DNA binding, organic cyclic compound binding* or *phosphatase activity*.

**Figure 3.**
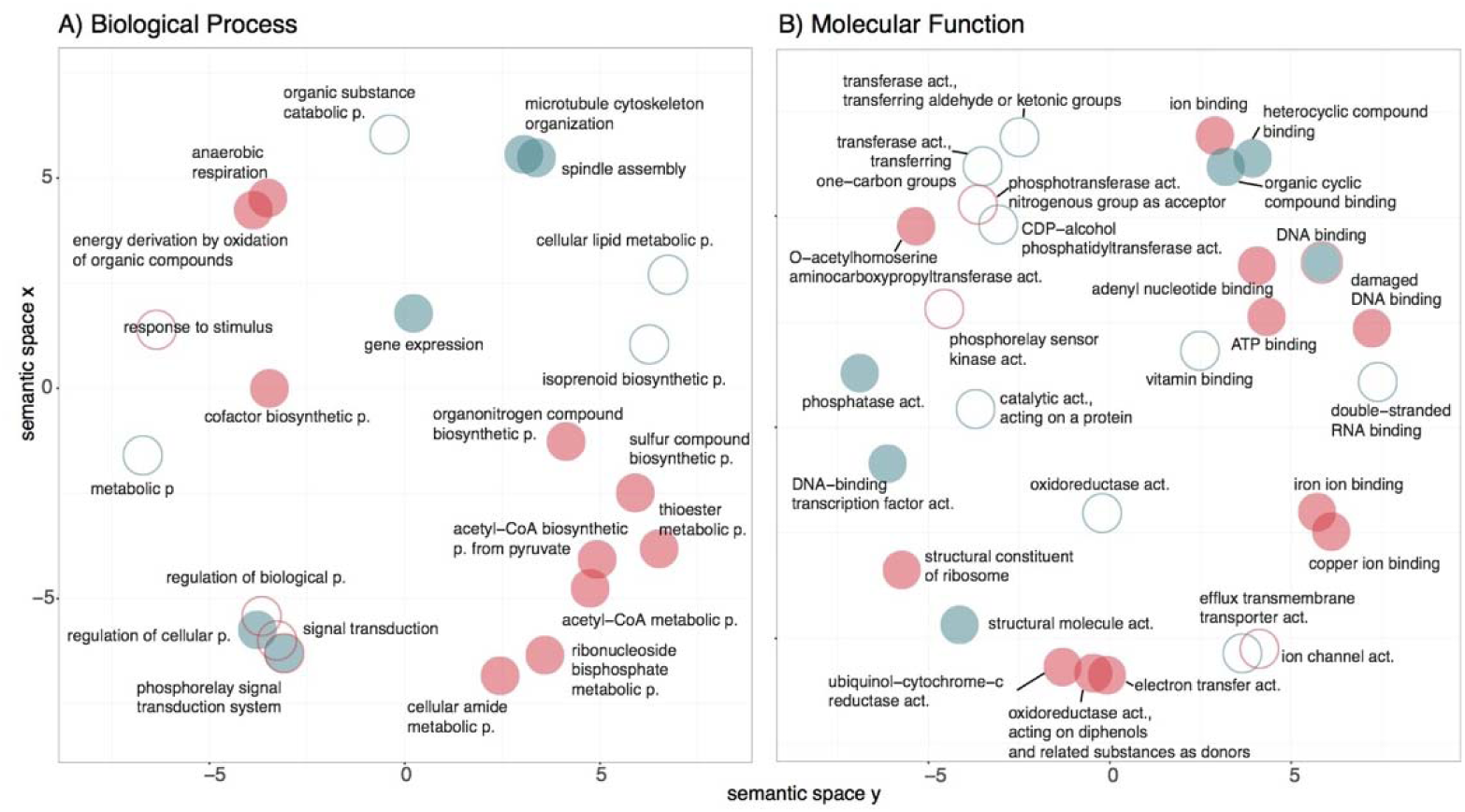
Gene Ontology (GO) enrichment analysis revealed metal-specific, over-represented terms ordered by semantic similarities in *Nitratiruptor* sp. SB155-2 cells when treated with cadmium or copper, relative to controls. A multidimensional scatterplot was created using the tool REVIGO (http://revigo.irb.hr/) and results were exported to the R environment, where they were plotted using the package, ggplot2. The following settings were used: semantic similarity: 0.7 (small), semantic similarity measure: SimRel. Open circles indicate GO terms associated with downregulated genes while solid circles indicate upregulation for *Biological Processes* (A, C) and *Molecular Function* (B, D).

In order to understand which KEGG pathways were enriched among DEGs, the hypergeometric test, Kegga, was used. A total of 16 and 12 signaling pathways were enriched following Cd and Cu exposure, respectively (p>0.05) (Supplemental Table 6). *Flagellar assembly, ribosome, RNA degradation* and *bacterial chemotaxis* were the pathways enriched among the upregulated gene sets following Cd exposure. The dominance of flagella formation in Cd treated samples vs. controls was further examined by negative staining by using TEM. While the proportion of flagellum found in control samples accounted for approximately half of the cells, in Cd treated samples more than 75% of the cells were found with flagellum (Supplemental Figure 1). As a result of Cu exposure, pathways related to *ribosome, microbial metabolism in diverse environments, sulfur metabolism, RNA degradation, nitrogen metabolism* and *oxidative phosphorylation* were over-represented. The log2FoldChange of the DEGs corresponding to some over-represented KEGG pathways are summarized in Table 2.

**Table 2.** Differentially expressed genes involved in A) Sulfur metabolism, B) Cysteine and methionine and C) Chemotaxis and flagella formation and expression ratios (in log2FoldChange) of *Nitratiruptor* sp. SB155-2 cells following cadmium (0.1 mM) or copper (0.05 mM) exposure.

|  | Gene ID | log2FoldChange |  | Gene name | Annotation |
| --- | --- | --- | --- | --- | --- |
|  |  | Cd | Cu |  |  |
| A) Sulfur metabolism |  |  |  |  |  |
| 1 | NIS_RS00170 |  | 1.82 |  | NAD(P)/FAD-dependent oxidoreductase |
| 2 | NIS_RS00180 |  | 2.31 | *soxY* | thiosulfate oxidation carrier protein SoxY |
| 3 | NIS_RS00185 |  | 1.57 | *soxZ* | thiosulfate oxidation carrier complex protein SoxZ |
| 4 | NIS_RS00795 |  | 2.45 |  | FAD-dependent oxidoreductase |
| 5 | NIS_RS00855 |  | 1.84 |  | NAD(P)/FAD-dependent oxidoreductase |
| 6 | NIS_RS01780 |  | 1.48 |  | FAD-dependent oxidoreductase |
| 7 | NIS_RS02255 |  | 1.9 |  | bifunctional oligoribonuclease/PAP phosphatase NrnA |
| 8 | NIS_RS09715 |  | 1.72 | *soxB* | thiosulfohydrolase SoxB |
| 9 | NIS_RS09720 |  | 1.94 | *soxA* | sulfur oxidation c-type cytochrome SoxA |
| 10 | NIS_RS09725 |  | 1.39 | *soxZ* | thiosulfate oxidation carrier complex protein SoxZ |
| B) Cysteine and methionine |  |  |  |  |  |
| 1 | NIS_RS03035 |  | 1.38 | *metH* | methionine synthase |
| 2 | NIS_RS06840 |  | 2.27 |  | O-acetylhomoserine aminocarboxypropyltransferase/cysteine synthase |
| 3 | NIS_RS06845 | 0.86* | 2.1 |  | O-acetylhomoserine aminocarboxypropyltransferase/cysteine synthase |
| 4 | NIS_RS08135 |  | 1.24 |  | S-adenosylmethionine decarboxylase proenzyme |
| 5 | NIS_RS08270 |  | 1.38 |  | homoserine dehydrogenase |
| 6 | NIS_RS08520 |  | 1.49 | *luxS* | S-ribosylhomocysteine lyase |
| C) Chemotaxis and flagella formation |  |  |  |  |  |
| 1 | NIS_RS01505 | 1.07 |  | *cheB* | chemotaxis-specific protein-glutamate methyltransferase CheB |
| 2 | NIS_RS03445 | 1.12 |  | *motB* | flagellar motor protein MotB |
| 3 | NIS_RS03500 | 1.23 |  | *motB* | flagellar motor protein MotB |
| 4 | NIS_RS05300 | 1.1 |  |  | MotA/TolQ/ExbB proton channel family protein |
| 5 | NIS_RS05305 | 1.34 |  |  | flagellar basal body-associated FliL family protein |
| 6 | NIS_RS05310 | 1.2 |  |  | flagellin |
| 7 | NIS_RS09980 | 1.4 |  | *flgL* | flagellar hook-associated protein FlgL |

### 2.5. STEM observation and presence of pPLGs granules

*Nitratiruptor* sp. B155-2 thin sections were observed using a transmission electron microscope in scanning mode (STEM) to investigate morphological differences under metal treatments vs. controls. No visible differences in cell morphology were observed in regard to the cytosol or cell wall (Supplemental Figure 3). Electron-dense round granules in the cytoplasm were confirmed under all treatments (Supplemental Figure 3-4) with moderately different numbers and sizes, depending on the treatment applied (Supplemental Figure 4). Figure 4 shows the spectral profiles resulting from energy-dispersive X-ray spectroscopy (EDX) square mapping at different locations (i.e., cell cytosol, matrix background and pPLG) of cells cultured with and without metal. Chemical analysis of intracellular granules confirmed enrichment with phosphorus and calcium, identifying them as polyphosphate-like granules (pPLGs) (Supplemental Figure 2, Figure 4-5). Relative composition analysis by EDX confirmed that pPLG phosphorus and calcium levels were similar under all treatments tested (Figure 5).

**Figure 4.**
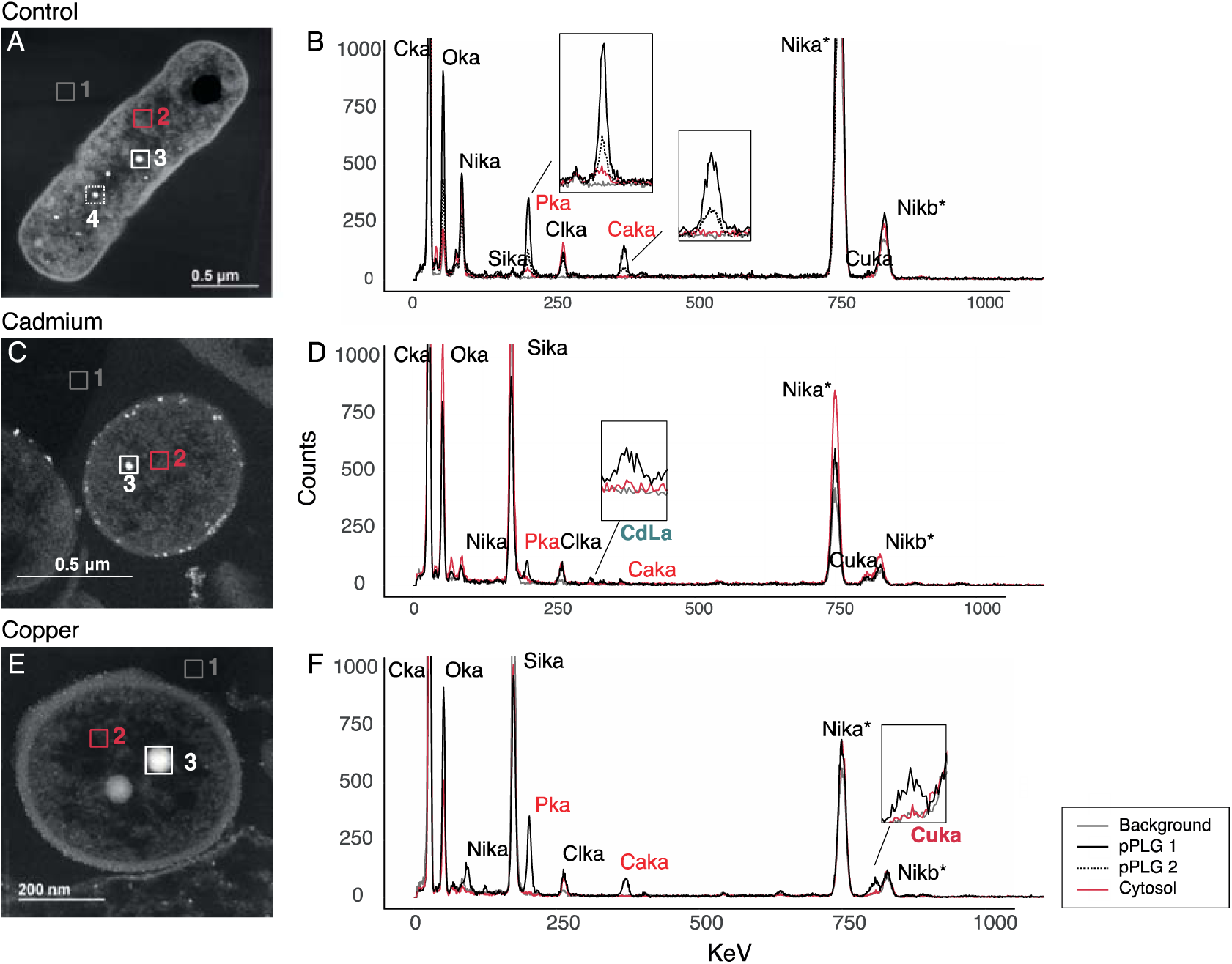
Scanning transmission electron microscopy (STEM) micrographs demonstrated the presence of polyphosphate-like granules (pPLGs) under all treatments (A. control, B. cadmium 0.1 mM and C. copper 0.05mM) while energy dispersive spectroscopy (EDX) analysis showed that pPLG elemental composition is metal-specific in *Nitratiruptor* sp. SB155-2. In each STEM micrograph, a square area mapping intracellular polyphosphate-like granules (pPLG), cytosol, and background was selected for elemental composition analysis and the resulting spectra are shown for the various treatments. Line color and style represent different mapping locations in the cell. A total of 10 individual cells per treatment were analyzed and produced similar spectral patterns. *Note that the high number of counts of Ni comes from the Ni grid supporting the sample.

**Figure 5.**
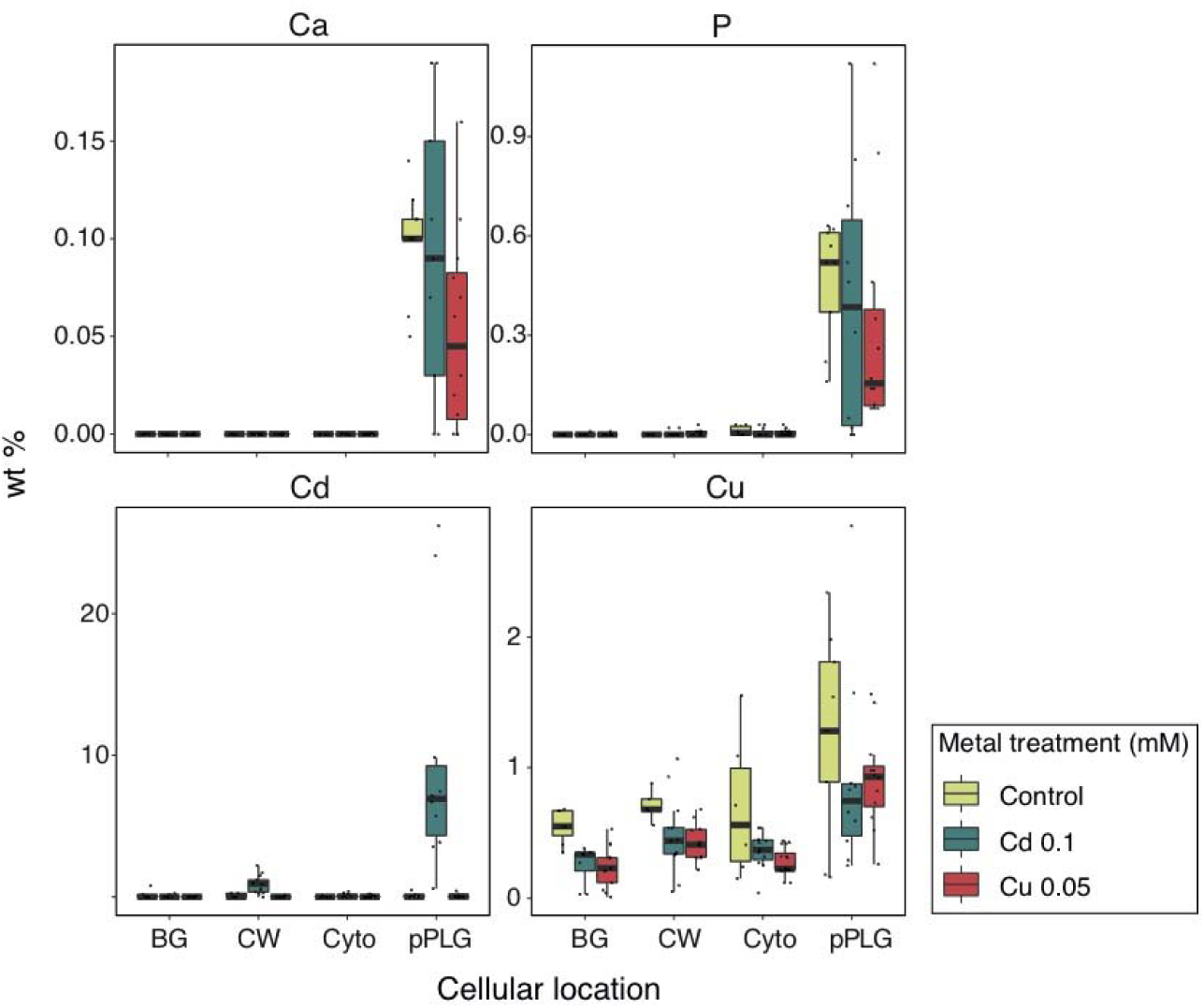
Cadmium treatment, but not copper, caused metal enrichment in polyphosphate-like granules in *Nitratiruptor* sp. SB155-2 cultures. Relative concentrations were compared at different cellular locations in *Nitratiruptor* sp. SB155-2 cells cultured in MMJS artificial seawater control (yellow) or with addition of Cd (0.1 mM) (blue) and Cu (0.05 mM) (red) for 24 h. Concentrations were estimated by area mapping using energy dispersive spectroscopy (EDX) and are shown in weight percent of dry mass (% wt). A total of 10 individual cells per treatment were analyzed as replicates. BG= Background, CW=cell wall, Cyto=cytoplasm, pPLG= polyphosphate-like granules.

### 2.6. Metal localization by EDX analysis

EDX analysis revealed the presence of Cd in pPLGs of cells grown under Cd exposure (Figure 4-5, Supplemental Figure 2). No Cd was detected in pPLGs of control cells or in those treated with Cu. Cd concentrations were higher in pPLG granules than in the cell wall, which also showed Cd-enriched particles (Figure 4-5, and Supplemental Figure 2). By contrast, measurable levels of Cu were found at different cell locations under all treatments tested, i.e., cytosol, cell wall, matrix background, and pPLG. No significant differences in Cu content were found in pPLGs between different treatments, because concentrations were highly variable. Additionally, using STEM, electron-dense particles were also observed attached to the outer membrane surface of cells following Cd exposure. Long-term mapping with EDX confirmed that the main elements included were Cd and S (Figure 6).

**Figure 6.**
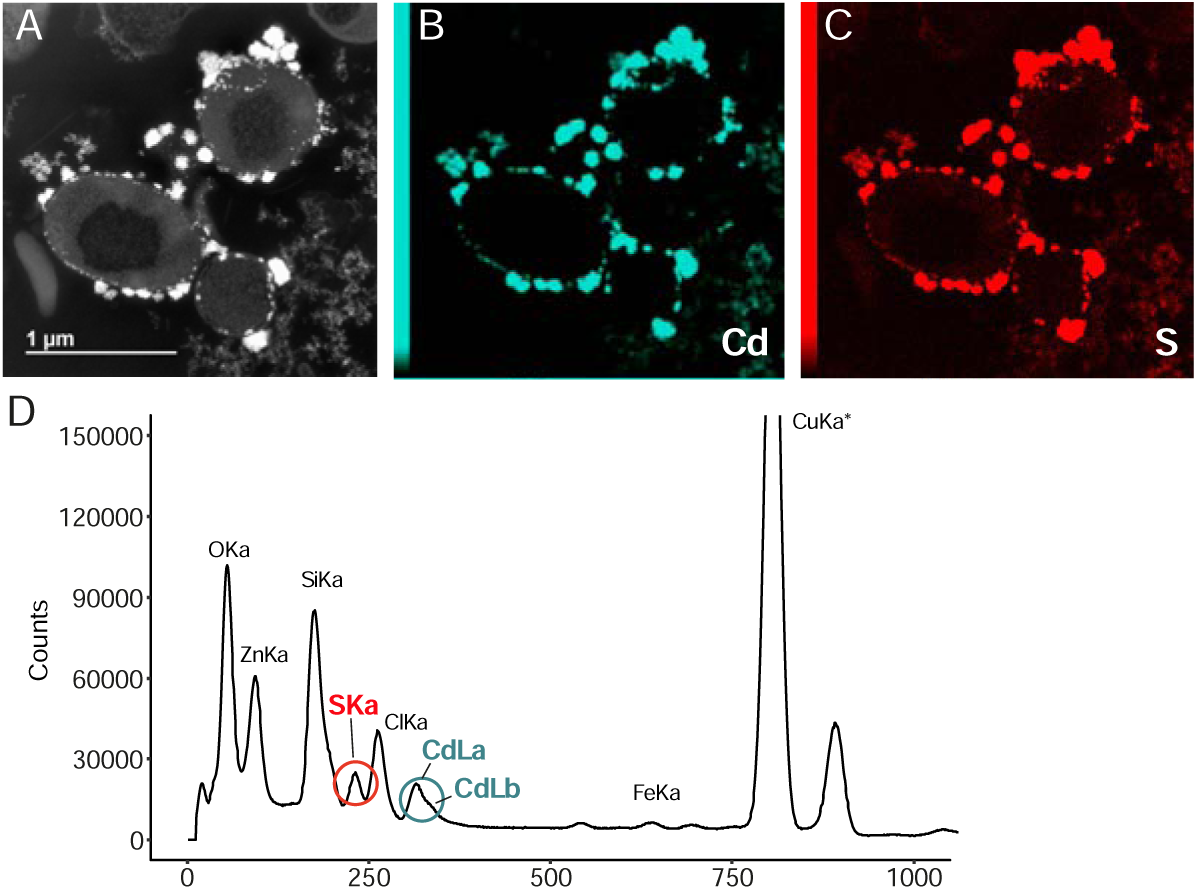
Cadmium treatment (0.1 mM, 24 hours) induced formation of CdS particles attached to the outer cell membrane in *Nitratiruptor* sp. SB155-2 cells. 15-h energy dispersive spectroscopy (EDX) mapping was carried out to determine the elemental composition of extracellular particles found in Cd-treated samples. Panel A-C shows the STEM micrograph, Cd and S localization in the area, respectively. D shows the energy dispersive spectrum analysis of the whole micrograph, indicating where the peaks for Cd and S are located. *Note that a high number of counts of Cu come from the Cu grid supporting the sample.

## 3. Discussion

The Epsilonproteobacteria is one of the predominant bacterial taxa inhabiting deep-sea hydrothermal vent ecosystems. A large body of literature has documented their contributions to vent biogeochemical processes, such as metabolic pathways to fix inorganic carbon, reduce nitrogen, or unique sulfur metabolism (Campbell *et al*., 2006; Yamamoto and Takai, 2011; Akerman *et al*., 2013; Vetriani *et al*., 2014). Nonetheless, adaptations to their metal-rich niche remain little-studied. This investigation examined the whole genome transcriptome of the deep-sea hydrothermal vent epsilonproteobacterium, *Nitratiruptor* sp. SB155-2, following exposure to Cd and Cu. In addition, high-throughput microscopy was applied to evaluate the condition of cells after exposure and to localize metal ions at the sub-cellular level. Predictably, metal treatment exhibited toxicity at the selected concentrations by suppressing the growth of *Nitratiruptor* sp. SB155-2 (Figure 1A) and inducing distinguishable transcriptomic responses to Cd and Cu. When so treated, 385 (19.3%) and 629 (31.6%) of genes were differentially expressed (Figure 1B,C-2). A more notable response triggered by Cu stress might be related to the essential functions this metal serves in biological activity. Cu is required for cell viability, but it is also highly toxic at relatively low concentrations in the cytoplasm, as opposed to nickel, zinc, manganese, requiring multiple layers of regulatory and protein-coding pathways to ensure cell homeostasis (Rademacher and Masepohl, 2012; Arguello *et al*., 2013; Bondarczuk and Piotrowska-Seget, 2013). The response to Cu appears more complex than to the non-bioessential Cd, including a larger number of high-affinity metal transporting systems, as well as oxidative stress-responsive genes (Supplemental Table 3-4). Additionally, sulfur, cysteine, and methionine metabolic pathways were upregulated only under Cu stress, while chemotaxis and flagella formation were upregulated only under Cd stress (Table 2, Figure 3). The various patterns in transcriptomic responses are consistent with our observations using electron microscopy and EDX. While pPLGs were present regardless of culture conditions, their role in metal sequestration varied, depending on the metal (Figure 4). While the Cd content of pPLGs was relatively high, a possible sequestration strategy for Cd, the role of pPLGs under Cu stress is still not clear. In some cells, EDX analysis of pPLGs showed relatively higher levels of Cu than in control or Cd-treated cells, while in others there were no visible differences between treatments (Figure 5). Additionally, Cd exposure triggered formation of CdS precipitates (Figure 6) as a Cd detoxification strategy. Here, we shed new light on Cd- and Cu-responsive molecular mechanisms of a deep-sea hydrothermal vent epsilonproteobacterium.

### 3.1. Differentially expressed genes common to Cd and Cu stress

Bacteria have developed efficient metal-specific adaptations (Hu *et al*., 2005; Lu *et al*., 2017; Villagrasa *et al*., 2021), and accordingly, *Nitatiruptor* sp. SB155-2’s transcriptomic profiles differ markedly when responding to Cd or Cu (Figure 2). Still, we found 27 genes (1.3% of the total genome) up-regulated in the presence of both metals (Table 1, Figure 2C), while another 50 (2.5%) were down-regulated in both (Supplemental Table 2, Figure 2C). Among these common DEGs, a large proportion are involved in metal efflux complexes (Nakagawa *et al*., 2007).

The first of the two gene clusters, encoding multicomponent transport systems, presumably correspond to the *ars* operon, a conserved arsenic (As) detoxification mechanism in gram-negative bacteria (Diorio *et al*., 1995). Arsenic resistance in bacteria is mediated through the arsRBC genes. arsR encodes an arsenic ion repressor that regulates expression of an arsenate reductase (arsC), involved in transforming As(V) into As(III) and an inner membrane-associated As(III) export system (arsB) responsible for export of As(III) from the cell (Moore *et al*., 2005). Our BLAST searches show that NIS_RS00140 and NIS_RS00160 are the most likely candidates for arsB and arsR, respectively, according to their genomic location and sequence similarity when compared to *E. coli* (strain K12) (E values 1e^-136^ and 2e^-16^ respectively). Previous studies found that transcription of arsR is less metal-specific than expected, given that it is also induced by other metals, such as zinc (Peng *et al*., 2018), Cd (Hu *et al*., 2005; Moore *et al*., 2005), antimony or bismuth (Hu *et al*., 2005; Moore *et al*., 2005); hence, initiating subsequent up-regulation of the *ars* operon. One possible explanation for induction of the *ars* operon following Cd exposure is the similarity of the metal-binding site of arsR to that of cadC, the metal-binding repressor for the *cad* operon and specific for Cd (Yoon and Silver, 1991), (Bairoch, 1993). Nevertheless, metal exposure experiments using *Cupriavidus metallidurans* CH34 revealed that other metals including Cu, cobalt, or nickel didn’t relieve the repression of arsR (Zhang *et al*., 2009), contrary to our observations for Cu. The second cluster encompasses several genes, including a permease annotated as SO_0444 family Cu/Zn efflux transporter, a cytochrome, and a TolC family protein. TolC family proteins are very versatile outer membrane channels (OMC) involved in export of compounds of different sizes and compositions (Tanabe et al., 2009; Turlin et al., 2014, Federici *et al*., 2004). This protein can function in combination with other efflux pumps, such as RND, ABC, or MFS, suggesting that this operon may also belong to one of these large families of efflux pumps, even though BLAST searches did not provide further hints about it. Importantly, genes displaying the highest log2FoldChange values for both elements, indicate that the SO_0444 family Cu/Zn efflux transporter seems to be central to Cd and Cu detoxification by *Nitratiruptor* sp. SB155-2, and a better characterization of this operon is needed in future studies. As its annotation suggests, this efflux transporter seems to be equally induced by Cu and Zn, as a result of nonspecific induction by Cd. Cadmium can replace zinc in Zn-requiring proteins, including efflux transport systems, as both elements share chemical and physical properties (Tang *et al*., 2014).

Excess metal content produces oxidative stress, threatening cell homeostasis. One of the main consequences is the loss of protein structure and function due to protein unfolding and misfolding (Imamoglu *et al*., 2020). In such a case, different chaperone systems, such as the well-known DnaK/DnaJ and GroES/GroEL systems, are rapidly and efficiently activated (Susin *et al*., 2006), and as expected, genes encoding both chaperone systems were up-regulated by both metals in *Nitratiruptor* sp. SB155-2. Also, glutamate 5-kinase is likely involved in stress alleviation. This enzyme catalyzes the transfer of a phosphate group to glutamate to form L-glutamate 5-phosphate and it is involved in synthesis of proline (Pérez-Arellano *et al*., 2006). This amino acid serves several protective functions as an osmoprotectant or scavenger of reactive oxygen species (ROS) (Meena *et al*., 2019); hence, it facilitates adaptation to environmental stress in bacteria (Siripornadulsil *et al*., 2002). For instance, (Al-Mailem *et al*., 2018) found that addition of proline to hypersaline soil containing heavy metals enhanced heavy metal tolerance in four halophilic and hydrocarbonoclastic bacteria, and proved useful for bioremediation. Additionally, protein oxidation, as a consequence of oxidative stress, results in their proteolytic degradation. Serine proteases have proven especially important for bacterial survival under multiple stress conditions (Zarzecka *et al*., 2019). The gene encoding a DegQ family serine endoprotease was highly up-regulated, indicating that it is strongly induced under metal stress, making this gene a putative biomarker for metal stress (Song *et al*., 2020).

### 3.2. Cadmium stress induces flagella formation and CdS production as major tolerance/detoxification mechanisms in *Nitratiruptor* sp. SB155-2

The genetic determinants, *czc* and *cad*, are major efflux systems that confer Cd resistance in bacteria. In addition to Cd, the protein complex, czcCBA, a member of the RND family, can also export cobalt, zinc, and nickel in a number of bacterial species, e.g., *Pseudomonas aeruginosa* (Ducret et al., 2020), *Escherichia coli* (Nies, 1995), *Ralstonia metallidurans* (Nies, 2003), and *Pseudamonas putida* (Peng et al., 2018). The *Nitratiruptor* sp SB155-2 genome includes one candidate gene for CzcB (i.e. NIS_RS03660) and another for CzcA (NIS_RS04945), while no gene candidates were found for CzcC. As a second line of defense, the P-type ATPase, CadA, also participates in Cd tolerance, being the primary determinant of Cd resistance in species like *Bacillus subtilis* (Moore *et al*., 2005) or *Staphylococcus aureus* (Nucifora et al., 1989). Genomic analysis of *Nitratiruptor* SB155-2 also confirmed the presence of a candidate gene encoding CadA P-type ATPase (i.e., NIS_RS07760). Unexpectedly, our transcriptomic results show that Cd stress did not up-regulate any of the genes encoding these two Cd-specific efflux transport systems (Supplemental Table 3). Instead, two ABC transporter permeases were differentially expressed exclusively upon exposure to Cd (Supplemental Table 3). The ATP-binding cassette (ABC) family, one of the largest transporter families, is involved in both import and export of substances, including toxic substances and can also contribute to metal homeostasis (Ma *et al*., 2009); however, few of these transporters likely contribute to Cd resistance.

According to the enrichment analysis (Figure 3, Table 2, Supplemental Table 6) chemotaxis and flagella formation were over-represented pathways under Cd stress. It seems reasonable to think that *Nitratiruptor* sp. SB155-2 cells can respond to toxic levels of Cd by activating flagella formation and chemical sensing capacity in order to find more favorable conditions. By activating these pathways, bacteria can move efficiently in response to steep, fast-changing gradient fluctuations in vent environments. In other gram-negative models, flagella formation under Cd stress causes opposing responses. Some species of bacteria lost mobility due to the loss of flagella (Siripornadulsil *et al*., 2014). In others, transcriptomic profiles show a clear up-regulation of genes encoding flagellar proteins as a primary response to Cd (Ma and Sun, 2021). Consistent with the latter, we confirmed that Cd induced formation of flagella, according to both transcriptomic evidence and negative staining in *Nitratiruptor* sp. SB155-2 cells (Supplemental Figure 1). The presence of this appendage would enable bacterial cells to find favorable environmental conditions for growth and survival in vent environments (Matilla and Krell, 2018).

Importantly, *Nitratiruptor* sp. SB155-2 cells produced CdS-enriched granules attached to the outer cell membrane only after Cd exposure (Figure 6). Production of CdS particles by bacteria growing under high Cd concentrations has been observed in other prokaryotes, including the deep-sea hydrothermal vent bacteria, *Idiomarina* sp. OT37-5b (Ma *et al*., 2020) and *Pseudoalteromonas* sp. MT33b (Ma and Sun, 2021). These authors found that methionine gamma-lyase was the main enzyme involved in formation of CdS particles through desulfurization of cysteine. The *Nitratiruptor* sp. SB155-2 genome does not possess a gene encoding a methionine gamma-lyase, but BLAST searches revealed two O-acetylhomoserine aminocarboxypropyltransferases (NIS_RS06840 and NIS_RS06845, not significant and padj < 0.05, respectively) under Cd stress. These were highly similar to the *Idiomarina* sp. OT37-5b methionine gamma-lyase, which may be involved in formation of CdS particles. Bacteria inhabiting vent environments could obtain cysteine from environmental sources, in the form of organo-sulfur molecules (OMS) (Ma *et al*., 2020). Organo-sulfur molecules contribute to sulfur cycling in hydrothermal vent environments, but bacterial uptake mechanisms are unknown (Wasmund *et al*., 2017).

### 3.3. *Nitratiruptor* sp. SB155-2 relies on efficient efflux and cysteine and methionine metabolism for Cu detoxification

The three main genetic determinants of Cu resistance in bacteria, *cus, cop*, and *cue* (Franke *et al*., 2003), are present in the *Nitratiruptor* sp SB155-2 genome. In bacteria, the RND complex, cusCFBA, is involved in direct export of periplasmic Cu out of the cell (Franke *et al*., 2001; Nies, 2003). While we found candidate genes for CusA and CusB, *Nitratiruptor* sp SB155-2 was devoid of candidate genes encoding CusC and CusF. Contrary to expectations, neither CusA nor CusB was differentially expressed under Cu stress, suggesting that *cus* may not be essential for Cu detoxification in these bacteria, at least, under the micro-aerobic conditions in this study. Activity of *cus* was dominant under anaerobic conditions (Outten *et al*., 2001), while the heavy metal translocator P-type ATPase, CopA, in combination with the periplasmic multicopper oxidase (CueO) usually operates under aerobic conditions (Outten *et al*., 2001). The main mechanism of the CopA-CueO export system involves exportation of Cu by CopA from the cytoplasm to the periplasm, where it is oxidized by CueO from Cu(I) to Cu(II) to protect periplasmic enzymes from copper-induced damage (Grass and Rensing, 2001). The *Nitratiruptor* sp. SB155-2 genome contains a CueO gene and sequence analysis revealed a P-type ATPase with great similarity to the *E. coli* CopA, a clear candidate for this P-type ATPase in Cu detoxification. Both CopA and CueO were significantly upregulated exclusively in samples treated with Cu (Supplemental Table 3-4), suggesting that the combination CopA-CueO is a major mechanism for Cu detoxification in *Nitratiruptor* sp SB155-2.

Periplasmic proteins enriched in the sulfur amino acids, methionine and cysteine, are able to bind Cu with high affinity. Other studies of gram-negative bacteria (Franke *et al*., 2003, Long et al., 2010, Lu *et al*., 2017) confirm that methionine residues are essential for Cu binding in the main Cu-specific transport systems. Crystal structure analysis of the inner membrane transporter, CusA, revealed four methionine pairs, in addition to the three methionine metal-binding sites located in the cleft of the periplasmic domain (Long *et al*., 2010). Because of this structure, CusA is capable of binding Cu directly from the cytosol and periplasm using methionine clusters (Su *et al*., 2011). Similarly, CopA, a methionine-rich periplasmic protein, is able to bind up to 11 Cu atoms (Puig *et al*., 2002). Other examples include periplasmic proteins, PcoC or CopC, methionine-rich proteins able to bind both Cu(I) and Cu(II) (Roberts *et al*., 2002; Wernimont *et al*., 2003; Bondarczuk and Piotrowska-Seget, 2013; Lawton *et al*., 2016). Consistent with these results, KEGG and GO enrichment analysis revealed that genes involved in sulfur, methionine, and cysteine metabolism pathways are over-represented among upregulated genes of *Nitratiruptor* sp. SB155-2 (Table 2, Figure 3, Supplemental Table 5-6), indicating that sulfur plays a central role in its Cu resistance. Other than its incorporation into periplasmic transport proteins, methionine is required for synthesis of glutathione, a universal antioxidant in response to heavy metal-induced oxidative stress (Stewart *et al*., 2020). Our results suggest that glutathione serves a fundamental function in Cu stress, since glutathione synthase is exclusively upregulated in Cu-treated samples (Supplemental Table 4).

### 3.4. pPLGs serve different functions under Cd and Cu stress

According to previous studies, two contrasting strategies have been described for the function of pPLGs in the presence of toxic metals: 1) the number of pPLGs per cell increases in order to sequester excess metal (Keasling and Hupf, 1996; Villagrasa *et al*., 2021); 2) pPLGs are hydrolysed in order to transport metal ions toward the periplasmic space bound to inorganic phosphate (Alvarez and Jerez, 2004; Seufferheld *et al*., 2008). Our results suggest that *Nitratiruptor* sp. SB155-2 adopts the first strategy for metal sequestration. Unlike eukaryotes, bacteria do not have discrete cellular compartments or organelles; thus, metal-binding metabolites, as well as storage and sequestration mechanisms, are crucial in order to keep metal concentrations under control (Chandrangsu *et al*., 2017). Among many other roles, pPLGs are involved in metal chelation, especially in bacteria that do not harbor superoxide dismutase (SOD), a highly conserved enzyme for detoxification of superoxide anion (Docampo, 2006). The *Nitratiruptor sp*. SB155-2 genome analysis revealed an absence of SOD (Nakagawa *et al*., 2007), implying that pPLGs serve as the primary sequestration mechanism in these bacteria. However, in *Nitratiruptor* sp. SB155-2 the function of pPLGs in sequestration may be somewhat Cd-specific when compared to Cu exposure. The number of pPLGs per cell is moderately higher than in controls, but Cd is highly concentrated in these granules. On the other hand, Cu seems to trigger formation of more and relatively larger pPLGs per cell, but the concentration of Cu in those granules is variable. (Figure 4-5, Supplemental Figure 2). While the reasons for an increased number of pPLGs under Cu stress remain unknown, its relatively low concentration compared to Cd, may be due to Cu’s contribution to various biological processes, as well as to the presence of efficient efflux mechanisms, which closely regulate intracellular Cu concentrations. Additionally, the number and size of pPLGs in this study should be interpreted with caution, since more precise microscopic techniques should be used to evaluate these features accurately, such as with FIB-SEMS, since 2D micrographs can only provide an incomplete picture of these 3D structures. Mobilization of pPLGs in bacteria is performed by the enzymes, polyphosphate kinase (PPK) and exopolyphosphatase (PPX), which synthesize and hydrolyze pPLGs, respectively. The *Nitratiruptor* sp. SB155-2 genome possesses two PPKs and two candidate PPXs. Surprisingly, neither PPK nor PPX was differentially expressed under conditions tested in this study. The lack of differential expression for these genes could be a result of a mismatch in the sampling timing for RNA extractions vs expression time required for these genes. Previous studies on the metal-resistant archaeon, *Metallosphaera sedula*, showed that a transcriptional shift of PPX occurred only 30 min after Cu exposure, and after 3h, PPX levels were again similar to those of controls (Rivero *et al*., 2018). In order to fully understand the transcriptional shift of these enzymes in *Nitratiruptor* sp. SB155-2 following metal exposure, future characterization should include a fine-scale time-course experiment to study expression level dynamics of these genes. In addition to pPLGs and CdS, high-resolution STEM enabled identification of Cd particles at the periphery of the cells (Figure 5), likely located in the periplasmic space. Together with pPLGs, the periplasmic space of gram-negative bacteria can be used as an intracellular compartment providing a storage or detoxification area to keep metal ions out of the cytosol (Ma *et al*., 2009).

Finally, even though pPLGs and other subcellular structures are well known in bacteria, application of modern high-resolution microscopic techniques in combination with high-throughput sequencing approaches can provide novel details regarding the timing and specific roles of these structures under metal stress. Future studies may benefit from these technologies to better understand detailed mechanisms of these structures regarding metal tolerance in microbes associated with deep-sea hydrothermal ecosystems. Prokaryotes inhabiting metal-enriched niches will reveal new metabolic capabilities and metal detoxification solutions (Antwis *et al*., 2017). These mechanisms could contribute significantly to new approaches to environmental restoration and remediation.

## 4. Experimental Procedures

### 4.1. Bacterial strain growth conditions

*Nitratiruptor* sp. SB155-2 was kindly provided by Satoshi Nakagawa (Kyoto University). Strain SB155-2 was maintained axenically in MMJS artificial seawater (Nunoura *et al*., 2008) containing NaNO_3_ and NaHCO_3_ (1g/L) at 55°C and a pH of 6.7, without shaking, as described in *et al*.. Glass bottles (100-mL) (Schott Duran, Germany) containing 30 mL of medium were used throughout the study. To initiate each culture, 0.5 mL of the inoculum (10^7^-10^8^ cells/mL) were added to 100-mL bottles. Tubes and bottles were closed with rubber stoppers and the head space gas was replaced with H_2_/CO_2_ (80:20) at a gas pressure of 0.2 MPa.

### 4.2. Heavy metal treatments

To test metal tolerance, *Nitratiruptor* SB155-B responses to Cd and Cu were evaluated at different concentrations (from 0.01 to 0.5 mM). Stock solutions of these elements were prepared by dissolving the respective salt (i.e., CdCl_2_ ·2.5 H_2_O and CuSO_4_ ·5 H_2_O for Cd and Cu treatments, respectively) in MilliQ water. To prepare working metal solutions, required volumes were added to the cultures by filtration through 0.22-µm membrane filters (Millipore Corp., United States).

#### Growth enumeration

Cell growth was studied under different treatments (Cd: 0.05, 0.1 and 0.5 mM; Cu: 0.01, 0.05 and 0.1 mM, n=4) and control cultures in order to determine appropriate sublethal concentrations. An aliquot of 1 mL was collected every 12 h for 5 days. Cell density was determined using standard flow cytometry (Brussaard, 2004; De Corte *et al*., 2012) with minor modifications. Briefly, sample aliquots were fixed with glutaraldehyde (0.5% final concentration) and frozen at -80°C until quantification. After thawing on ice, the sample was stained with SYBR Green I (Molecular Probes, Invitrogen, Carlsbad, USA). Cell numbers were counted using an Accuri C6 flow cytometer (BD Biosciences, US) and fluorescence versus side scatter was plotted.

#### Incubation for STEM-EDX and RNA-seq analysis

Once the cultures reached the late-exponential phase, i.e., after 4 days, metals were included at a final concentration of 0.01mM for Cd and 0.05mM for Cu. Bottles were incubated for 24 h (n=3) for microscopy and 3h for RNA-seq analysis (n=4). After the incubation period, samples were immediately used for STEM-EDX preparation or rapidly filtered through 0.2-µm PTFE filters (Merck, Germany), flash frozen in liquid nitrogen, and stored at -80°C until RNA extraction.

### 4.3. Electron microscopy observation

#### STEM-EDX observation

After metal incubation, *Nitratiruptor* sp. SB155B cells were collected by centrifugation at 4000 rpm for 5 min at 4 °C. Fixation was carried out in 2.5% (v/v) glutaraldehyde in 0.1M sodium cacodylate buffer for 2 h in darkness at room temperature. Then, the supernatant was removed and samples were post-fixed with Osmium tetroxide (OsO_4_) for 1 h. Samples were dehydrated in an ethanol series (70%, 80%, 90%, 95% and 100%) and finally embedded in resin. Fixed samples were cut using a Leica UC6 microtome (Leica, Germany) with a diamond cutter (DiATOME, US) at a thickness of 100 nm. Finally, sections mounted on carbon-coated nickel grids (Nisshin-EM, Japan) were examined by STEM with a JEOL ARM-200F system at 80Kv with an angular annular dark field detector (30-120 mrad) using HR STEM-HAADF observation mode. The spot size was 6C for observing and 1C for EDX mapping.

For EDX detection, a JEOL SDD high-resolution EDX detector (100mm^2^ solid angle) was used. Elemental spectra were collected from at least two points per cell for a minimum of 30 cells per treatment. For characterization, total cell composition was mapped in at least at five cells per treatment for 5-15 h. Atomic composition of ratios and distribution were assessed using the software, JED-2300 Analysis Station.

#### TEM observation of flagella

In order to examine whether Cd-treated cultures produced larger proportions of flagella, negative staining evaluation was performed with TEM. Cells from Cd-treated and control samples (n=3) were gently collected by centrifugation at 2000 rpm for 4 minutes at 4°C, in order to not disrupt the flagella and a drop was collected and directly mounted on carbon-coated nickel grids. Samples were negatively stained with 1% uranyl acetate and observed under a JEM1230R electron microscope at an accelerating voltage of 100keV. More than 200 cells were photographed and examined for presence/absence of flagella.

### 4.4. Total RNA isolation, library preparation, and sequencing

Total RNA was extracted from the filters using ZR Fungal/Bacterial RNA kits (Zymo Research, US) following manufacturer instructions. Samples were treated with DNase I (Qiagen, Germany) for DNA removal. Following extraction, RNA quality and concentration were assessed on an Agilent 2100 Bioanalyzer and a Qubit 2.0 fluorometer, respectively. After quality was determined, 2 replicates for Cd and Cu treatments and 3 replicates for controls met the quality criteria to proceed with subsequent steps. Total rRNA was extracted from the samples using Ribo-zero Magnetic kits (Illumina, US). For library preparation, NEBNext® Ultra Directional RNA Library Prep Kits for Illumina® (Illumina, US) were used following manufacturer instructions. Quality control of prepared libraries was determined with an Agilent 2100 Bioanalyzer, and after library normalization, cDNA libraries were sequenced on an Illumina NovaSeqTM 6000 sequencing system with a 2 × 150-bp pair-end read length protocol.

### 4.5. RNA-seq data analyses

The resulting FASTA files were processed using the Nextflow pipeline nfcore/rnaseq (version 3.1) mainly with standard settings (Ewels *et al*., 2020). However, the few changes made to the settings are summarized as follows: i) strandedness of the library was set as reverse in the input file; ii) Hisat2 was the aligner selected; iii) in order to remove adaptors and low-quality sequences, the Trim Galore clipped length was changed to 15 bp. Reads were mapped to the reference sequence *Nitratiruptor* sp. SB155-2 (GenBank: Assembly: GCA_000010325.1). Gene counts for each sample were extracted from StringTie results using the python script, prepDE.py and imported into the R statistical environment for further analysis. In order to identify potential outliers and major sources of variation, hierarchical clustering heatmap and principal component analysis were performed after RLD normalization. Differential gene expression analysis between metal-treated cultures was performed with the DESeq function in the Bioconductor package, DESeq2 (Love *et al*., 2014). Genes that were considered differentially expressed with a False Discovery Rate (FDR) adjusted p-value (padj) less than 0.05 and a change of at least 2-fold (log2FoldChange= 1) were considered statistically significant. Gene Ontology (GO) information was obtained using Blast2Go software (version 5.2.5). Identification of GO terms enriched among differentially expressed genes was carried out with the hypergeometric test in the R package, GOstats (Falcon and Gentleman 2007). Differentially enriched GO terms were visualized in semantic similarity-based scatterplots using REVIGO (http://revigo.irb.hr/). Identification of enriched KEGG pathways was further investigated by applying the Kegga function in the R package, edgeR (Robinson et al. 2010). Both GO terms and KEGG pathways were considered significantly enriched with a p-value less than 0.05. Sequencing data have been deposited in the NCBI Sequencing Read Archive under accession PRJNA746661

## 5. Acknowledgements

The authors thank Margaret Mars Brisbin and Charles Plessy for their advice on bioinformatic analysis and constructive feedback, and Yuko Hasegawa for providing helpful technical assistance. We thank Steven D. Aird for English editing and helpful feedback. We are grateful for help and support provided by the Scientific Computing and Data Analysis section of the Research Support Division at the Okinawa Institute of Science and Technology. This work was supported by funding from the Marine Biophysics Unit (OIST) of the Okinawa Institute of Science and Technology. TN was partially supported by JSPS KAKENHI Grant Number JP19H05684 within JP19H05679 (Post-Koch Ecology).

## References

Akerman, N.H., Butterfield, D.A., and Huber, J.A. (2013) Phylogenetic diversity and functional gene patterns of sulfur-oxidizing subseafloor Epsilonproteobacteria in diffuse hydrothermal vent fluids. Front Microbiol 4: 185.

Al-Mailem, D.M., Eliyas, M., and Radwan, S.S. (2018) Ferric Sulfate and Proline Enhance Heavy-Metal Tolerance of Halophilic/Halotolerant Soil Microorganisms and Their Bioremediation Potential for Spilled-Oil Under Multiple Stresses. Front Microbiol 9: 394.

Alvarez, S. and Jerez, C.A. (2004) Copper ions stimulate polyphosphate degradation and phosphate efflux in Acidithiobacillus ferrooxidans. Appl Environ Microbiol 70: 5177–5182.

Antwis, R.E., Griffiths, S.M., Harrison, X.A., Aranega-Bou, P., Arce, A., Bettridge, A.S., et al. (2017) Fifty important research questions in microbial ecology. FEMS Microbiol Ecol 93.:

Arguello, J., Raimunda, D., and Padilla-Benavides, T. (2013) Mechanisms of copper homeostasis in bacteria. Front Cell Infect Microbiol 3: 73.

Bairoch, A. (1993) A possible mechanism for metal-ion induced DNA-protein dissociation in a family of prokaryotic transcriptional regulators. Nucleic Acids Res 21: 2515.

Ben Fekih, I., Zhang, C., Li, Y.P., Zhao, Y., Alwathnani, H.A., Saquib, Q., et al. (2018) Distribution of Arsenic Resistance Genes in Prokaryotes. Front Microbiol 9: 2473.

Bondarczuk, K. and Piotrowska-Seget, Z. (2013) Molecular basis of active copper resistance mechanisms in Gram-negative bacteria. Cell Biol Toxicol 29: 397–405.

Brussaard, C.P.D. (2004) Optimization of procedures for counting viruses by flow cytometry. Appl Environ Microbiol 70: 1506–1513.

Campbell, B.J., Engel, A.S., Porter, M.L., and Takai, K. (2006) The versatile ε-proteobacteria: key players in sulphidic habitats. Nat Rev Microbiol 4: 458–468.

Chandrangsu, P., Rensing, C., and Helmann, J.D. (2017) Metal homeostasis and resistance in bacteria. Nat Rev Microbiol 15: 338–350.

Chen, H.-Y., Huh, C.-A., Chang, N.-Y., and Chen, J.-C. (2000) Sources and Distribution of Heavy Metals in East China Sea Surface Sediments. Chemistry and Ecology 17: 227–238.

Cobine, P.A., Pierrel, F., and Winge, D.R. (2006) Copper trafficking to the mitochondrion and assembly of copper metalloenzymes. Biochimica et Biophysica Acta (BBA) - Molecular Cell Research 1763: 759–772.

De Corte, D., Sintes, E., Yokokawa, T., Reinthaler, T., and Herndl, G.J. (2012) Links between viruses and prokaryotes throughout the water column along a North Atlantic latitudinal transect. ISME J 6: 1566–1577.

Diorio, C., Cai, J., Marmor, J., Shinder, R., and DuBow, M.S. (1995) An Escherichia coli chromosomal ars operon homolog is functional in arsenic detoxification and is conserved in gram-negative bacteria. J Bacteriol 177: 2050–2056.

Docampo, R. (2006) Acidocalcisomes and Polyphosphate Granules. In Inclusions in Prokaryotes. Shively, J.M. (ed). Berlin, Heidelberg: Springer Berlin Heidelberg, pp. 53–70.

Ducret, V., Gonzalez, M.R., Leoni, S., Valentini, M., and Perron, K. (2020) The CzcCBA Efflux System Requires the CadA P-Type ATPase for Timely Expression Upon Zinc Excess in Pseudomonas aeruginosa. Front Microbiol 11: 911.

Ewels, P.A., Peltzer, A., Fillinger, S., Patel, H., Alneberg, J., Wilm, A., et al. (2020) The nf-core framework for community-curated bioinformatics pipelines. Nat Biotechnol 38: 276–278.

Fan, B., Grass, G., Rensing, C., and Rosen, B.P. (2001) Escherichia coli CopA N-Terminal Cys(X)2Cys Motifs Are Not Required for Copper Resistance or Transport. Biochem Biophys Res Commun 286: 414–418.

Federici, L., Walas, F., and Luisi, B. (2004) The structure and mechanism of the TolC outer membrane transport protein. Curr Sci 87: 190–196.

Franke, S., Grass, G., and Nies, D.H. (2001) The product of the ybdE gene of the Escherichia coli chromosome is involved in detoxification of silver ions. Microbiology 147: 965–972.

Franke, S., Grass, G., Rensing, C., and Nies, D.H. (2003) Molecular analysis of the copper-transporting efflux system CusCFBA of Escherichia coli. J Bacteriol 185: 3804–3812.

Fukui, T., Atomi, H., Kanai, T., Matsumi, R., Fujiwara, S., and Imanaka, T. (2005) Complete genome sequence of the hyperthermophilic archaeon Thermococcus kodakaraensis KOD1 and comparison with Pyrococcus genomes. Genome Res 15: 352–363.

Gort, A.S., Ferber, D.M., and Imlay, J.A. (1999) The regulation and role of the periplasmic copper, zinc superoxide dismutase of Escherichia coli. Mol Microbiol 32: 179–191.

Grass, G. and Rensing, C. (2001) CueO is a multi-copper oxidase that confers copper tolerance in Escherichia coli. Biochem Biophys Res Commun 286: 902–908.

Hu, P., Brodie, E.L., Suzuki, Y., McAdams, H.H., and Andersen, G.L. (2005) Whole-genome transcriptional analysis of heavy metal stresses in Caulobacter crescentus. J Bacteriol 187: 8437–8449.

Imamoglu, R., Balchin, D., Hayer-Hartl, M., and Hartl, F.U. (2020) Bacterial Hsp70 resolves misfolded states and accelerates productive folding of a multi-domain protein. Nat Commun 11: 365.

Jiang, Z., Jiang, L., Zhang, L., Su, M., Tian, D., Wang, T., et al. (2020) Contrasting the Pb (II) and Cd (II) tolerance of Enterobacter sp. via its cellular stress responses. Environ Microbiol 22: 1507–1516.

Keasling, J.D. and Hupf, G.A. (1996) Genetic manipulation of polyphosphate metabolism affects cadmium tolerance in Escherichia coli. Appl Environ Microbiol 62: 743–746.

Kimber, R.L., Bagshaw, H., Smith, K., Buchanan, D.M., Coker, V.S., Cavet, J.S., and Lloyd, J.R. (2020) Biomineralization of Cu2S Nanoparticles by Geobacter sulfurreducens. Appl Environ Microbiol 86.:

Lagorce, A., Fourçans, A., Dutertre, M., Bouyssiere, B., Zivanovic, Y., and Confalonieri, F. (2012) Genome-wide transcriptional response of the archaeon Thermococcus gammatolerans to cadmium. PLoS One 7: e41935.

Lawton, T.J., Kenney, G.E., Hurley, J.D., and Rosenzweig, A.C. (2016) The CopC Family: Structural and Bioinformatic Insights into a Diverse Group of Periplasmic Copper Binding Proteins. Biochemistry 55: 2278–2290.

Long, F., Su, C.-C., Zimmermann, M.T., Boyken, S.E., Rajashankar, K.R., Jernigan, R.L., and Yu, E.W. (2010) Crystal structures of the CusA efflux pump suggest methionine-mediated metal transport. Nature 467: 484–488.

Lopez-Garcia, P., Duperron, S., Philippot, P., Foriel, J., Susini, J., and Moreira, D. (2003) Bacterial diversity in hydrothermal sediment and epsilonproteobacterial dominance in experimental microcolonizers at the Mid-Atlantic Ridge. Environmental Microbiology 5: 961–976.

Love, M.I., Huber, W., and Anders, S. (2014) Moderated estimation of fold change and dispersion for RNA-seq data with DESeq2. Genome Biol 15: 550.

Lu, M., Jiao, S., Gao, E., Song, X., Li, Z., Hao, X., et al. (2017) Transcriptome Response to Heavy Metals in Sinorhizobium meliloti CCNWSX0020 Reveals New Metal Resistance Determinants That Also Promote Bioremediation by Medicago lupulina in Metal-Contaminated Soil. Appl Environ Microbiol 83.:

Ma, N., Sha, Z., and Sun, C. (2020) Formation of cadmium sulfide nanoparticles mediates cadmium resistance and light utilization of the deep-sea bacterium Idiomarina sp. OT37-5b. Environ Microbiol.

Ma, N. and Sun, C. (2021) Cadmium sulfide nanoparticle biomineralization and biofilm formation mediate cadmium resistance of the deep-sea bacterium Pseudoalteromonas sp. MT33b. Environ Microbiol Rep.

Matilla, M.A. and Krell, T. (2018) The effect of bacterial chemotaxis on host infection and pathogenicity. FEMS Microbiol Rev 42.:

Ma, Z., Jacobsen, F.E., and Giedroc, D.P. (2009) Coordination chemistry of bacterial metal transport and sensing. Chem Rev 109: 4644–4681.

Meena, M., Divyanshu, K., Kumar, S., Swapnil, P., Zehra, A., Shukla, V., et al. (2019) Regulation of L-proline biosynthesis, signal transduction, transport, accumulation and its vital role in plants during variable environmental conditions. Heliyon 5: e02952.

Moore, C.M., Gaballa, A., Hui, M., Ye, R.W., and Helmann, J.D. (2005) Genetic and physiological responses of Bacillus subtilis to metal ion stress. Mol Microbiol 57: 27–40.

Nakagawa, S., Takaki, Y., Shimamura, S., Reysenbach, A.-L., Takai, K., and Horikoshi, K. (2007) Deep-sea vent epsilon-proteobacterial genomes provide insights into emergence of pathogens. Proc Natl Acad Sci U S A 104: 12146–12150.

Naz, N., Young, H.K., Ahmed, N., and Gadd, G.M. (2005) Cadmium accumulation and DNA homology with metal resistance genes in sulfate-reducing bacteria. Appl Environ Microbiol 71: 4610–4618.

Nies, D.H. (2003) Efflux-mediated heavy metal resistance in prokaryotes. FEMS Microbiol Rev 27: 313–339.

Nies, D.H. (1995) The cobalt, zinc, and cadmium efflux system CzcABC from Alcaligenes eutrophus functions as a cation-proton antiporter in Escherichia coli. J Bacteriol 177: 2707–2712.

Nucifora, G., Chu, L., Misra, T.K., and Silver, S. (1989) Cadmium resistance from Staphylococcus aureus plasmid pI258 cadA gene results from a cadmium-efflux ATPase. Proc Natl Acad Sci U S A 86: 3544–3548.

Nunoura, T., Miyazaki, M., Suzuki, Y., Takai, K., and Horikoshi, K. (2008) Hydrogenivirga okinawensis sp. nov., a thermophilic sulfur-oxidizing chemolithoautotroph isolated from a deep-sea hydrothermal field, Southern Okinawa Trough. Int J Syst Evol Microbiol 58: 676–681.

Outten, F.W., Huffman, D.L., Hale, J.A., and O’Halloran, T.V. (2001) The Independent cue and cusSystems Confer Copper Tolerance during Aerobic and Anaerobic Growth inEscherichia coli *. J Biol Chem 276: 30670–30677.

Outten, F.W., Outten, C.E., Hale, J., and O’Halloran, T.V. (2000) Transcriptional activation of an Escherichia coli copper efflux regulon by the chromosomal MerR homologue, cueR. J Biol Chem 275: 31024–31029.

Peng, J., Miao, L., Chen, X., and Liu, P. (2018) Comparative Transcriptome Analysis of Pseudomonas putida KT2440 Revealed Its Response Mechanisms to Elevated Levels of Zinc Stress. Front Microbiol 9: 1669.

Pérez-Arellano, I., Rubio, V., and Cervera, J. (2006) Mapping active site residues in glutamate-5-kinase. The substrate glutamate and the feed-back inhibitor proline bind at overlapping sites. FEBS Lett 580: 6247–6253.

Puig, S., Rees, E.M., and Thiele, D.J. (2002) The ABCDs of periplasmic copper trafficking. Structure 10: 1292–1295.

Rademacher, C. and Masepohl, B. (2012) Copper-responsive gene regulation in bacteria. Microbiology 158: 2451–2464.

Reyes-Caballero, H., Campanello, G.C., and Giedroc, D.P. (2011) Metalloregulatory proteins: metal selectivity and allosteric switching. Biophys Chem 156: 103–114.

Reysenbach, A.L., Banta, A.B., Boone, D.R., Cary, S.C., and Luther, G.W. (2000) Microbial essentials at hydrothermal vents. Nature 404: 835.

Rivero, M., Torres-Paris, C., Muñoz, R., Cabrera, R., Navarro, C.A., and Jerez, C.A. (2018) Inorganic Polyphosphate, Exopolyphosphatase, and Pho84-Like Transporters May Be Involved in Copper Resistance in Metallosphaera sedula DSM 5348T. Archaea 2018: 5251061.

Roberts, S.A., Weichsel, A., Grass, G., Thakali, K., Hazzard, J.T., Tollin, G., et al. (2002) Crystal structure and electron transfer kinetics of CueO, a multicopper oxidase required for copper homeostasis in Escherichia coli. Proc Natl Acad Sci U S A 99: 2766–2771.

Seufferheld, M.J., Alvarez, H.M., and Farias, M.E. (2008) Role of polyphosphates in microbial adaptation to extreme environments. Appl Environ Microbiol 74: 5867–5874.

Siripornadulsil, S., Thanwisai, L., and Siripornadulsil, W. (2014) Changes in the proteome of the cadmium-tolerant bacteria Cupriavidus taiwanensis KKU2500-3 in response to cadmium toxicity. Can J Microbiol 60: 121–131.

Siripornadulsil, S., Traina, S., Verma, D.P.S., and Sayre, R.T. (2002) Molecular mechanisms of proline-mediated tolerance to toxic heavy metals in transgenic microalgae. Plant Cell 14: 2837–2847.

Song, W.-S., Kim, S.-M., Jo, S.-H., Lee, J.-S., Jeon, H.-J., Ko, B.J., et al. (2020) Multi-omics characterization of the osmotic stress resistance and protease activities of the halophilic bacterium Pseudoalteromonas phenolica in response to salt stress. RSC Advances 10: 23792–23800.

Stewart, L.J., Ong, C.-L.Y., Zhang, M.M., Brouwer, S., McIntyre, L., Davies, M.R., et al. (2020) Role of Glutathione in Buffering Excess Intracellular Copper in Streptococcus pyogenes. MBio 11.:

Su, C.-C., Long, F., and Yu, E.W. (2011) The Cus efflux system removes toxic ions via a methionine shuttle. Protein Sci 20: 6–18.

Susin, M.F., Baldini, R.L., Gueiros-Filho, F., and Gomes, S.L. (2006) GroES/GroEL and DnaK/DnaJ have distinct roles in stress responses and during cell cycle progression in Caulobacter crescentus. J Bacteriol 188: 8044–8053.

Takai, K., Inagaki, F., Nakagawa, S., Hirayama, H., Nunoura, T., Sako, Y., et al. (2003) Isolation and phylogenetic diversity of members of previously uncultivated ε-Proteobacteria in deep-sea hydrothermal fields. FEMS Microbiol Lett 218: 167–174.

Takai, K. and Nakamura, K. (2010) Compositional, Physiological and Metabolic Variability in Microbial Communities Associated with Geochemically Diverse, Deep-Sea Hydrothermal Vent Fluids. In Geomicrobiology: Molecular and Environmental Perspective. Barton, L.L., Mandl, M., and Loy, A. (eds). Dordrecht: Springer Netherlands, pp. 251–283.

Tang, L., Qiu, R., Tang, Y., and Wang, S. (2014) Cadmium–zinc exchange and their binary relationship in the structure of Zn-related proteins: a mini review. Metallomics 6: 1313–1323.

Vetriani, C., Voordeckers, J.W., Crespo-Medina, M., O’Brien, C.E., Giovannelli, D., and Lutz, R.A. (2014) Deep-sea hydrothermal vent Epsilonproteobacteria encode a conserved and widespread nitrate reduction pathway (Nap). ISME J 8: 1510–1521.

Villagrasa, E., Egea, R., Ferrer-Miralles, N., and Solé, A. (2020) Genomic and biotechnological insights on stress-linked polyphosphate production induced by chromium(III) in Ochrobactrum anthropi DE2010. World Journal of Microbiology and Biotechnology 36.:

Villagrasa, E., Palet, C., López-Gómez, I., Gutiérrez, D., Esteve, I., Sánchez-Chardi, A., and Solé, A. (2021) Cellular strategies against metal exposure and metal localization patterns linked to phosphorus pathways in Ochrobactrum anthropi DE2010. J Hazard Mater 402: 123808.

Wang, B., Zeng, C., Chu, K.H., Wu, D., Yip, H.Y., Ye, L., and Wong, P.K. (2017) Enhanced biological hydrogen production fromEscherichia coliwith surface precipitated cadmium sulfide nanoparticles. Adv Energy Mater 7: 1700611.

Wasmund, K., Mußmann, M., and Loy, A. (2017) The life sulfuric: microbial ecology of sulfur cycling in marine sediments. Environ Microbiol Rep 9: 323–344.

Wernimont, A.K., Huffman, D.L., Finney, L.A., Demeler, B., O’Halloran, T.V., and Rosenzweig, A.C. (2003) Crystal structure and dimerization equilibria of PcoC, a methionine-rich copper resistance protein from Escherichia coli. J Biol Inorg Chem 8: 185–194.

Yamamoto, M. and Takai, K. (2011) Sulfur Metabolisms in Epsilon-and Gamma-Proteobacteria in Deep-Sea Hydrothermal Fields. Frontiers in Microbiology 2.:

Yang, Z., Lu, L., Berard, V.F., He, Q., Kiely, C.J., Berger, B.W., and McIntosh, S. (2015) Biomanufacturing of CdS quantum dots. Green Chem 17: 3775–3782.

Yoon, K.P. and Silver, S. (1991) A second gene in the Staphylococcus aureus cadA cadmium resistance determinant of plasmid pI258. J Bacteriol 173: 7636–7642.

Zarzecka, U., Harrer, A., Zawilak-Pawlik, A., Skorko-Glonek, J., and Backert, S. (2019) Chaperone activity of serine protease HtrA of Helicobacter pylori as a crucial survival factor under stress conditions. Cell Commun Signal 17: 161.

Zhang, K., Xue, Y., Xu, H., and Yao, Y. (2019) Lead removal by phosphate solubilizing bacteria isolated from soil through biomineralization. Chemosphere 224: 272–279.

Zhang, Y.-B., Monchy, S., Greenberg, B., Mergeay, M., Gang, O., Taghavi, S., and van der Lelie, D. (2009) ArsR arsenic-resistance regulatory protein from Cupriavidus metallidurans CH34. Antonie Van Leeuwenhoek 96: 161–170.

