## Supplemental material for "Sequestration and efflux largely account for cadmium and copper resistance in the deep sea epsilonproteobacterium, *Nitratiruptor* sp. SB155-2"

**Supplemental Table 1.** Sequencing statistics for each sample before and after trimming and mapping into the reference genome.

**Supplemental Table 2.** Common differentially down-regulated genes of *Nitratiruptor* SB155-2 following cadmium (0.1 mM) or copper (0.05) exposure, showing log2FoldChange and gene description.

**Supplemental Table 3.** List of over-represented GO terms among differentially expressed genes by application of the hypergeometric test in the R package, GOstats. Terms are classified as *Molecular Function* (MF), *Biological Processes* (BP) and *Cellular Component* (CC).

**Supplemental Table 4.** List of over-represented KEGG pathways among differentially expressed genes by application of the hypergeometric test, Kegga.

**Supplemental Figure 1.** Negatively stained *Nitratiruptor* sp. SB155-2 cells showing the presence of flagellum (A) and absence (B). Percentage of cells with and without flagellum under Cd (n=209) and control cultures (n=234).

**Supplemental Figure 2.** Presence of polyphosphate-like granules (pPLGs) in *Nitratiruptor* sp. SB155-2 cells under different treatments

**Supplemental Figure 3.** Long-term energy dispersive spectroscopy (EDX) mapping of *Nitratiruptor* sp. SB155-2 cells with addition of Cd (0.1 mM), Cu (0.05mM) and controls.

**Supplemental Figure 4.** A-C) Micrographs showing the presence of polyphosphate-like granules (pPLGs) *Nitratiruptor* sp. SB155-2 cells under different treatments. D) Barplot showing the number of pPLGs found per cell under different treatments. E) Boxplot showing the mean size distribution of pPLGs

| **Sample** | **Replicate** | **Sequenced read pairs** | **Total trimmed read pairs** | **Total mapped** |
| --- | --- | --- | --- | --- |
| Control | 1 | 5313463 | 5276268 (99.3%) | 4959693 (94.00%) |
|  | 2 | 4974318 | 4934523 (99.20%) | 4579238 (92.80%) |
|  | 3 | 6594539 | 6541783 (99.20%) | 5992273 (91.6%) |
| Cd | 1 | 5236304 | 5189177 (99.10%) | 4748097 (91.5%) |
|  | 2 | 4878787 | 4829999 (99%) | 3762569 (77.9%) |
| Cu | 1 | 5592347 | 5553201 (99.30%) | 4831284 (87%) |
|  | 2 | 5111202 | 5080535 (99.40) | 4155877 (81.80%) |

Table 1

Table 2

|  | **Ensgenes** | **log2FoldChange** | | **Gene name** | **Annotation** |
| --- | --- | --- | --- | --- | --- |
|  |  | **Cd** | **Cu** |  |  |
| 1 | NIS_RS00095 | -2.63 | -1.29 |  | ABC transporter ATP-binding protein |
| 2 | NIS_RS00100 | -1.53 | -1.19 |  | ABC transporter permease subunit |
| 3 | NIS_RS00610 | -2.76 | -2.33 |  | heme-binding domain-containing protein |
| 4 | NIS_RS00615 | -1.34 | -1.56 |  | HAMP domain-containing histidine kinase |
| 5 | NIS_RS00620 | -1.34 | -2.25 |  | response regulator transcription factor |
| 6 | NIS_RS01010 | -1.48 | -1.38 |  | right-handed parallel beta-helix repeat-containing protein |
| 7 | NIS_RS01520 | -1.06 | -2.3 | *CheW* | chemotaxis protein CheW |
| 8 | NIS_RS01645 | -2.47 | -1.18 |  | NADH-quinone oxidoreductase subunit M |
| 9 | NIS_RS01935 | -1.6 | -1.22 |  | hypothetical protein |
| 10 | NIS_RS02485 | -1.57 | -2.06 |  | hypothetical protein |
| 11 | NIS_RS03380 | -1.24 | -1.49 | *FlhF* | flagellar biosynthesis protein FlhF |
| 12 | NIS_RS03675 | -1.86 | -2.33 |  | DUF2905 domain-containing protein |
| 13 | NIS_RS03925 | -1.22 | -1.36 | *HyfE* | Ni-Fe hydrogenase, membrane subunit HyfE |
| 14 | NIS_RS04250 | -1.74 | -1.36 | *HisB* | imidazoleglycerol-phosphate dehydratase HisB |
| 15 | NIS_RS04255 | -2.17 | -1.35 |  | HAD-IIIA family hydrolase |
| 16 | NIS_RS04260 | -1.07 | -1.49 | *LptC* | LPS export ABC transporter periplasmic protein LptC |
| 17 | NIS_RS04315 | -1.39 | -1.36 |  | APC family permease |
| 18 | NIS_RS04730 | -1.8 | -1.3 |  | 1-acyl-sn-glycerol-3-phosphate acyltransferase |
| 19 | NIS_RS04805 | -1.67 | -1.06 |  | hypothetical protein |
| 20 | NIS_RS05025 | -1.11 | -1.05 |  | TIGR00730 family Rossman fold protein |
| 21 | NIS_RS05775 | -1.15 | -1.94 |  | PAS domain-containing protein |
| 22 | NIS_RS05945 | -2.46 | -2.12 | *LptF/LptG* | LptF/LptG family permease |
| 23 | NIS_RS06005 | -1.86 | -1.85 |  | hypothetical protein |
| 24 | NIS_RS06300 | -1.54 | -1.05 | *CvpA* | CvpA family protein |
| 25 | NIS_RS06925 | -1.65 | -2.1 |  | hypothetical protein |
| 26 | NIS_RS07070 | -2 | -1.85 |  | AEC family transporter |
| 27 | NIS_RS07270 | -1.5 | -1.3 |  | prepilin-type N-terminal cleavage/methylation domain-containing protein |
| 28 | NIS_RS07275 | -1.8 | -1.94 |  | hypothetical protein |
| 29 | NIS_RS07470 | -1.72 | -1.02 |  | glycosyl transferase |
| 30 | NIS_RS08365 | -1.83 | -1.34 |  | type II secretion system protein |
| 31 | NIS_RS09270 | -1.26 | -1.44 | *TrkH* | TrkH family potassium uptake protein |
| 32 | NIS_RS09355 | -1.42 | -1.23 |  | EAL domain-containing protein |
| 33 | NIS_RS09645 | -1.4 | -1.23 |  | 4Fe-4S binding protein |

Table 3

|  | **GO_id** | **Pvalue** | **OddsRatio** | **ExpCount** | **Count** | **Size** | **Term** | **Ont** | **DE** | **elem** |
| --- | --- | --- | --- | --- | --- | --- | --- | --- | --- | --- |
| 1 | GO:0051225 | 0.038095238 | Inf | 0.038095238 | 1 | 1 | spindle assembly | BP | Up | Cd |
| 2 | GO:0000226 | 0.038095238 | Inf | 0.038095238 | 1 | 1 | microtubule cytoskeleton organization | BP | Up | Cd |
| 3 | GO:0000160 | 0.03971599 | 5.778947368 | 0.838095238 | 3 | 22 | phosphorelay signal transduction system | BP | Up | Cd |
| 4 | GO:0050794 | 0.040873074 | 4.771428571 | 1.485714286 | 4 | 39 | regulation of cellular process | BP | Up | Cd |
| 5 | GO:0010467 | 0.044804608 | 5.46 | 0.876190476 | 3 | 23 | gene expression | BP | Up | Cd |
| 6 | GO:0044255 | 0.008143043 | 8.133333333 | 0.942857143 | 4 | 9 | cellular lipid metabolic process | BP | Down | Cd |
| 7 | GO:0008299 | 0.029554656 | 18.7 | 0.314285714 | 2 | 3 | isoprenoid biosynthetic process | BP | Down | Cd |
| 8 | GO:1901575 | 0.029554656 | 18.7 | 0.314285714 | 2 | 3 | organic substance catabolic process | BP | Down | Cd |
| 9 | GO:0008152 | 0.031363303 | 3.426229508 | 14.77142857 | 19 | 141 | metabolic process | BP | Down | Cd |
| 10 | GO:0015980 | 0.002812405 | 33.35294118 | 0.380952381 | 3 | 4 | energy derivation by oxidation of organic compounds | BP | Up | Cu |
| 11 | GO:0043603 | 0.006598193 | 16.58823529 | 0.476190476 | 3 | 5 | cellular amide metabolic process | BP | Up | Cu |
| 12 | GO:0009061 | 0.008658009 | Inf | 0.19047619 | 2 | 2 | anaerobic respiration | BP | Up | Cu |
| 13 | GO:0051188 | 0.02463525 | 5.027777778 | 1.238095238 | 4 | 13 | cofactor biosynthetic process | BP | Up | Cu |
| 14 | GO:1901566 | 0.039373602 | 3.111801242 | 2.761904762 | 6 | 29 | organonitrogen compound biosynthetic process | BP | Up | Cu |
| 15 | GO:0035383 | 0.046138644 | 10.44444444 | 0.380952381 | 2 | 4 | thioester metabolic process | BP | Up | Cu |
| 16 | GO:0071616 | 0.046138644 | 10.44444444 | 0.380952381 | 2 | 4 | acyl-CoA biosynthetic process | BP | Up | Cu |
| 17 | GO:0034032 | 0.046138644 | 10.44444444 | 0.380952381 | 2 | 4 | purine nucleoside bisphosphate metabolic process | BP | Up | Cu |
| 18 | GO:0006084 | 0.046138644 | 10.44444444 | 0.380952381 | 2 | 4 | acetyl-CoA metabolic process | BP | Up | Cu |
| 19 | GO:0006086 | 0.046138644 | 10.44444444 | 0.380952381 | 2 | 4 | acetyl-CoA biosynthetic process from pyruvate | BP | Up | Cu |
| 20 | GO:0033866 | 0.046138644 | 10.44444444 | 0.380952381 | 2 | 4 | nucleoside bisphosphate biosynthetic process | BP | Up | Cu |
| 21 | GO:0033875 | 0.046138644 | 10.44444444 | 0.380952381 | 2 | 4 | ribonucleoside bisphosphate metabolic process | BP | Up | Cu |
| 22 | GO:0044272 | 0.046138644 | 10.44444444 | 0.380952381 | 2 | 4 | sulfur compound biosynthetic process | BP | Up | Cu |
| 23 | GO:0050789 | 0.000531379 | 4.349794239 | 6.247619048 | 14 | 41 | regulation of biological process | BP | Down | Cu |
| 24 | GO:0007165 | 0.000997519 | 4.656084656 | 4.419047619 | 11 | 29 | signal transduction | BP | Down | Cu |
| 25 | GO:0050896 | 0.003739917 | 3.507692308 | 5.79047619 | 12 | 38 | response to stimulus | BP | Down | Cu |
| 26 | GO:0000160 | 0.008345449 | 3.904761905 | 3.352380952 | 8 | 22 | phosphorelay signal transduction system | BP | Down | Cu |
| 27 | GO:0003700 | 0.001138618 | 43.24137931 | 0.271714922 | 3 | 4 | DNA-binding transcription factor activity | MF | Up | Cd |
| 28 | GO:0003677 | 0.008544478 | 2.563041655 | 5.773942094 | 12 | 85 | DNA binding | MF | Up | Cd |
| 29 | GO:0005198 | 0.01083771 | 3.088534107 | 3.192650334 | 8 | 47 | structural molecule activity | MF | Up | Cd |
| 30 | GO:0016791 | 0.024926964 | 14.15254237 | 0.271714922 | 2 | 4 | phosphatase activity | MF | Up | Cd |
| 31 | GO:0097159 | 0.034011354 | 1.722176325 | 17.18596882 | 24 | 253 | organic cyclic compound binding | MF | Up | Cd |
| 32 | GO:1901363 | 0.034011354 | 1.722176325 | 17.18596882 | 24 | 253 | heterocyclic compound binding | MF | Up | Cd |
| 33 | GO:0015562 | 0.005676917 | 8.888888889 | 0.837416481 | 4 | 8 | efflux transmembrane transporter activity | MF | Down | Cd |
| 34 | GO:0017169 | 0.010852806 | Inf | 0.20935412 | 2 | 2 | CDP-alcohol phosphatidyltransferase activity | MF | Down | Cd |
| 35 | GO:0003725 | 0.010852806 | Inf | 0.20935412 | 2 | 2 | double-stranded RNA binding | MF | Down | Cd |
| 36 | GO:0019842 | 0.010852806 | Inf | 0.20935412 | 2 | 2 | vitamin binding | MF | Down | Cd |
| 37 | GO:0016491 | 0.027884775 | 1.832951945 | 11.51447661 | 18 | 110 | oxidoreductase activity | MF | Down | Cd |
| 38 | GO:0016744 | 0.030329716 | 17.45652174 | 0.31403118 | 2 | 3 | transferase activity, transferring aldehyde or ketonic groups | MF | Down | Cd |
| 39 | GO:0140096 | 0.042217326 | 2.19491256 | 4.815144766 | 9 | 46 | catalytic activity, acting on a protein | MF | Down | Cd |
| 40 | GO:0016741 | 0.049048313 | 2.95505618 | 2.093541203 | 5 | 20 | transferase activity, transferring one-carbon groups | MF | Down | Cd |
| 41 | GO:0003735 | 3.18E-06 | 4.585804133 | 8.667037862 | 22 | 43 | structural constituent of ribosome | MF | Up | Cu |
| 42 | GO:0016679 | 0.008080297 | Inf | 0.60467706 | 3 | 3 | oxidoreductase activity, acting on diphenols and related substances as donors | MF | Up | Cu |
| 43 | GO:0009055 | 0.018677549 | 2.866968326 | 4.395089286 | 9 | 22 | electron transfer activity | MF | Up | Cu |
| 44 | GO:0030554 | 0.026684009 | 1.769472666 | 14.91536748 | 22 | 74 | adenyl nucleotide binding | MF | Up | Cu |
| 45 | GO:0043167 | 0.031496288 | 1.501831502 | 31.84632517 | 41 | 158 | ion binding | MF | Up | Cu |
| 46 | GO:0005506 | 0.03765299 | 2.844252874 | 3.426503341 | 7 | 17 | iron ion binding | MF | Up | Cu |
| 47 | GO:0003684 | 0.040446626 | Inf | 0.40311804 | 2 | 2 | damaged DNA binding | MF | Up | Cu |
| 48 | GO:0005507 | 0.040446626 | Inf | 0.40311804 | 2 | 2 | copper ion binding | MF | Up | Cu |
| 49 | GO:0003961 | 0.040446626 | Inf | 0.40311804 | 2 | 2 | O-acetylhomoserine aminocarboxypropyltransferase activity | MF | Up | Cu |
| 50 | GO:0008121 | 0.040446626 | Inf | 0.40311804 | 2 | 2 | ubiquinol-cytochrome-c reductase activity | MF | Up | Cu |
| 51 | GO:0005524 | 0.042909839 | 1.678485577 | 14.71380846 | 21 | 73 | ATP binding | MF | Up | Cu |
| 52 | GO:0000155 | 0.030538717 | 4.192207792 | 1.262806236 | 4 | 14 | phosphorelay sensor kinase activity | MF | Down | Cu |
| 53 | GO:0016775 | 0.030538717 | 4.192207792 | 1.262806236 | 4 | 14 | phosphotransferase activity, nitrogenous group as acceptor | MF | Down | Cu |
| 54 | GO:0003677 | 0.033372394 | 1.978145425 | 7.667037862 | 13 | 85 | DNA binding | MF | Down | Cu |
| 55 | GO:0005216 | 0.042778904 | 10.3164557 | 0.360801782 | 2 | 4 | ion channel activity | MF | Down | Cu |

Table 4

| **ID** | **Pathway** | **N** | **DE** | **P.DE** |  |
| --- | --- | --- | --- | --- | --- |
| path:nis02040 | Flagellar assembly | 35 | 6 | 0.003655706 | Cdup |
| path:nis03010 | Ribosome | 62 | 7 | 0.017840985 | Cdup |
| path:nis03018 | RNA degradation | 13 | 3 | 0.018319804 | Cdup |
| path:nis02030 | Bacterial chemotaxis | 25 | 4 | 0.023398765 | Cdup |
| path:nis01100 | Metabolic pathways | 425 | 56 | 0.003857535 | Cddown |
| path:nis00190 | Oxidative phosphorylation | 39 | 10 | 0.004515164 | Cddown |
| path:nis00630 | Glyoxylate and dicarboxylate metabolism | 13 | 5 | 0.007254893 | Cddown |
| path:nis01230 | Biosynthesis of amino acids | 95 | 17 | 0.012862189 | Cddown |
| path:nis01200 | Carbon metabolism | 60 | 12 | 0.015900685 | Cddown |
| path:nis01110 | Biosynthesis of secondary metabolites | 215 | 31 | 0.019469795 | Cddown |
| path:nis00564 | Glycerophospholipid metabolism | 11 | 4 | 0.020680996 | Cddown |
| path:nis01120 | Microbial metabolism in diverse environments | 116 | 19 | 0.021838539 | Cddown |
| path:nis01503 | Cationic antimicrobial peptide (CAMP) resistance | 7 | 3 | 0.028291411 | Cddown |
| path:nis00680 | Methane metabolism | 19 | 5 | 0.039558018 | Cddown |
| path:nis02020 | Two-component system | 61 | 11 | 0.042954393 | Cddown |
| path:nis00260 | Glycine, serine and threonine metabolism | 14 | 4 | 0.049156712 | Cddown |
| path:nis03010 | Ribosome | 62 | 26 | 6.62E-06 | Cuup |
| path:nis01120 | Microbial metabolism in diverse environments | 116 | 36 | 0.000281001 | Cuup |
| path:nis00920 | Sulfur metabolism | 19 | 10 | 0.000727135 | Cuup |
| path:nis03018 | RNA degradation | 13 | 6 | 0.020240797 | Cuup |
| path:nis00910 | Nitrogen metabolism | 17 | 7 | 0.024422395 | Cuup |
| path:nis00190 | Oxidative phosphorylation | 39 | 12 | 0.041431999 | Cuup |
| path:nis02010 | ABC transporters | 22 | 9 | 2.40E-05 | Cudown |
| path:nis02040 | Flagellar assembly | 35 | 8 | 0.005894452 | Cudown |
| path:nis00564 | Glycerophospholipid metabolism | 11 | 4 | 0.009409603 | Cudown |
| path:nis02030 | Bacterial chemotaxis | 25 | 6 | 0.013505494 | Cudown |
| path:nis00561 | Glycerolipid metabolism | 4 | 2 | 0.037042089 | Cudown |
| path:nis04122 | Sulfur relay system | 10 | 3 | 0.043806577 | Cudown |

Figure 1

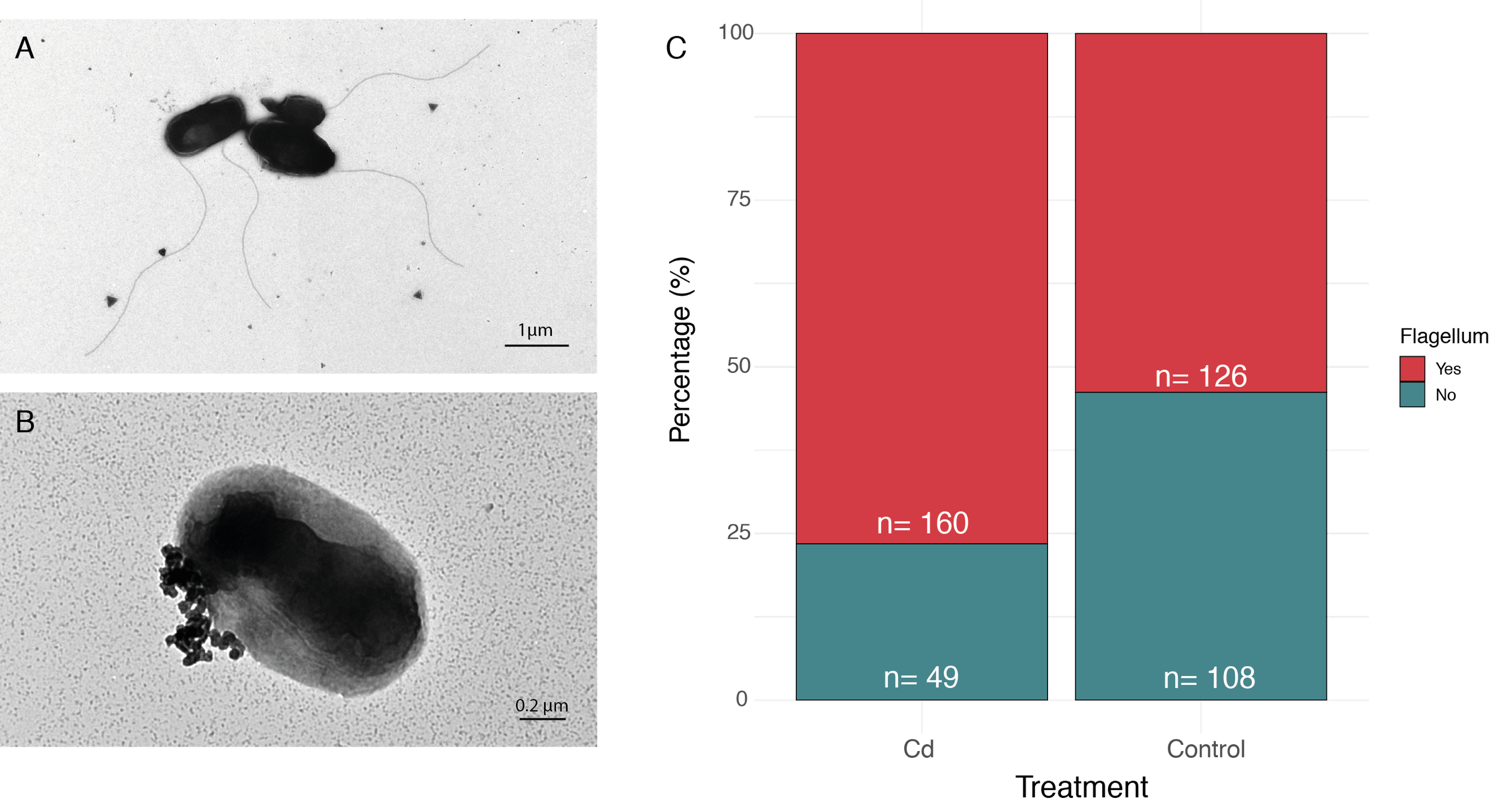

Figure 2

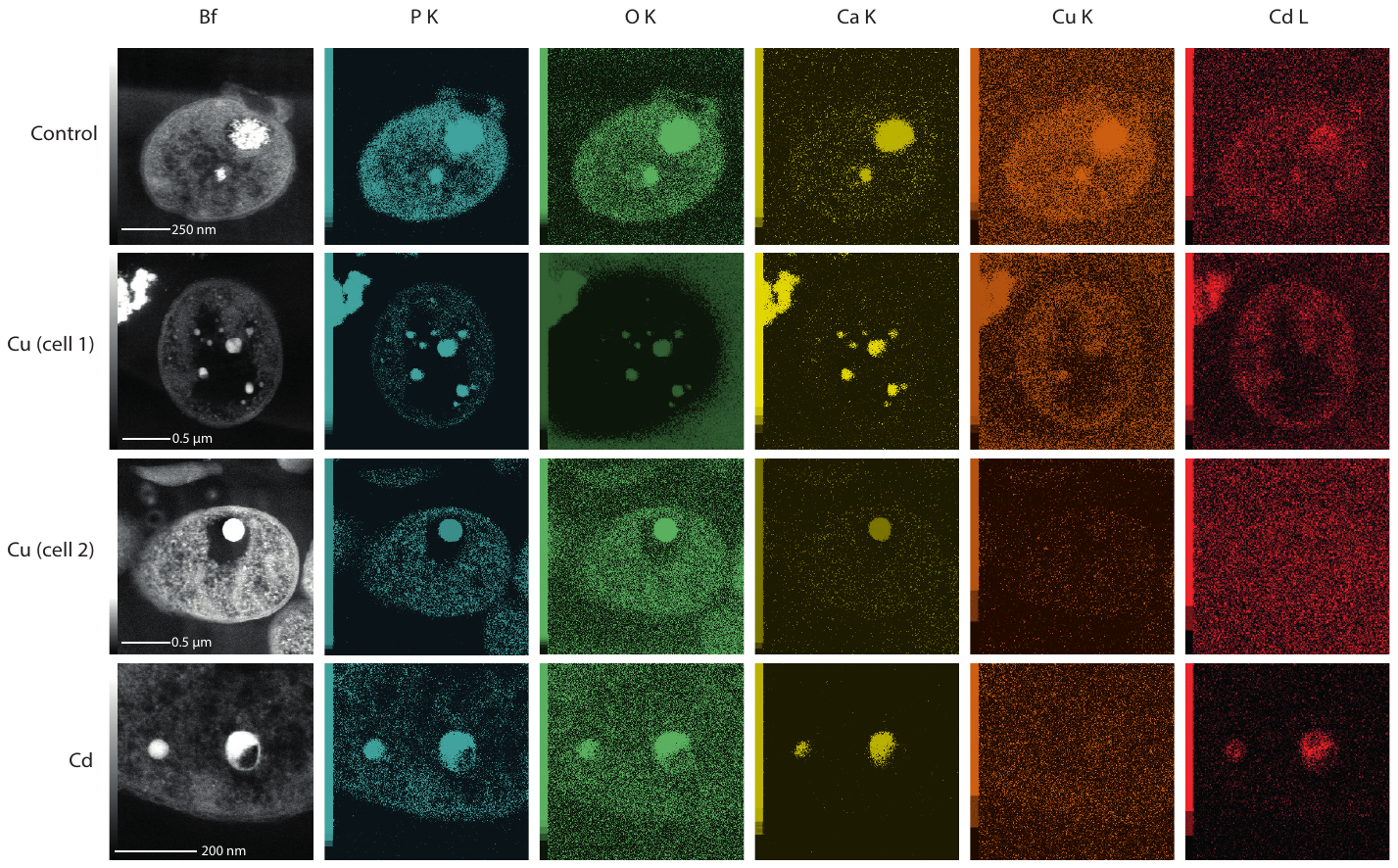

Figure 2

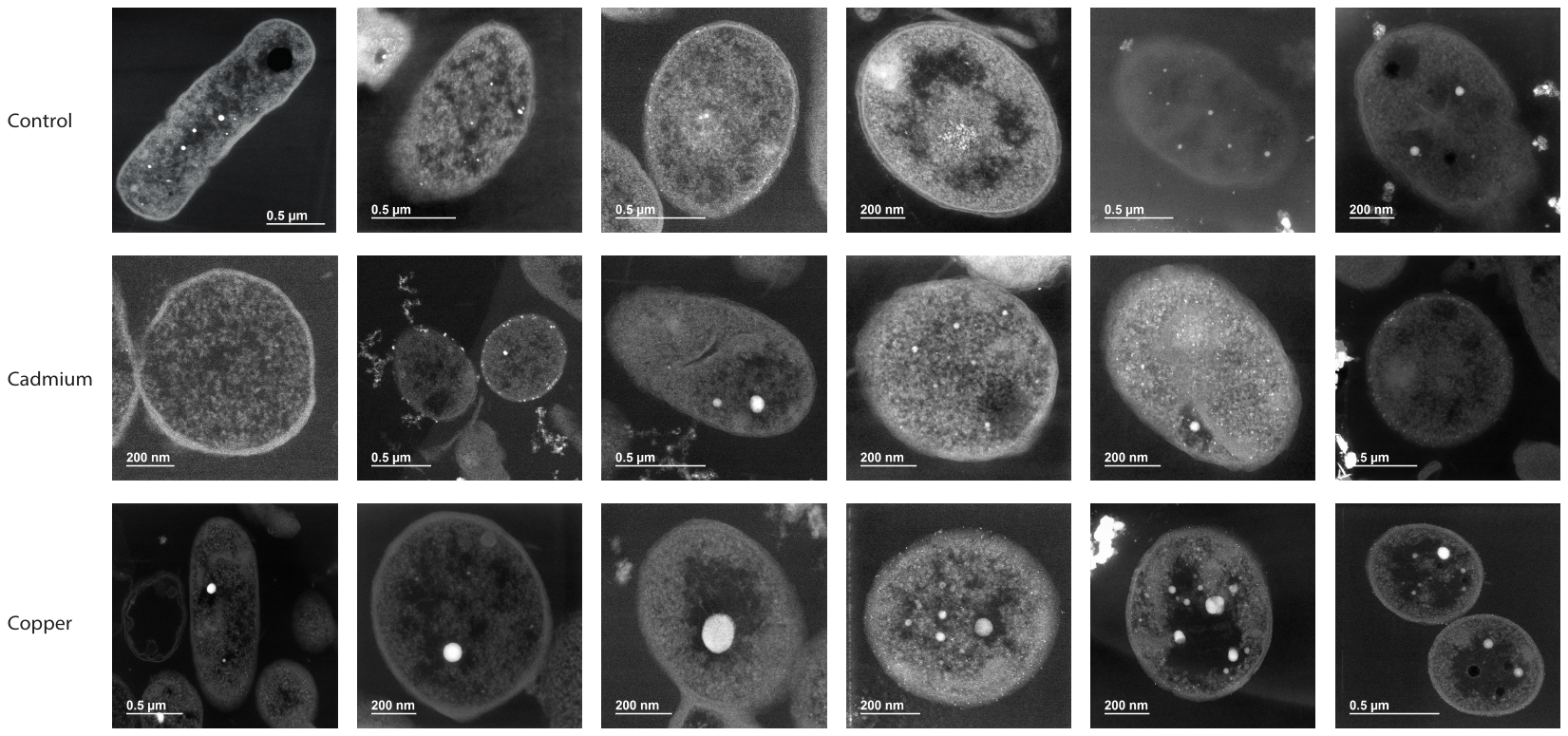

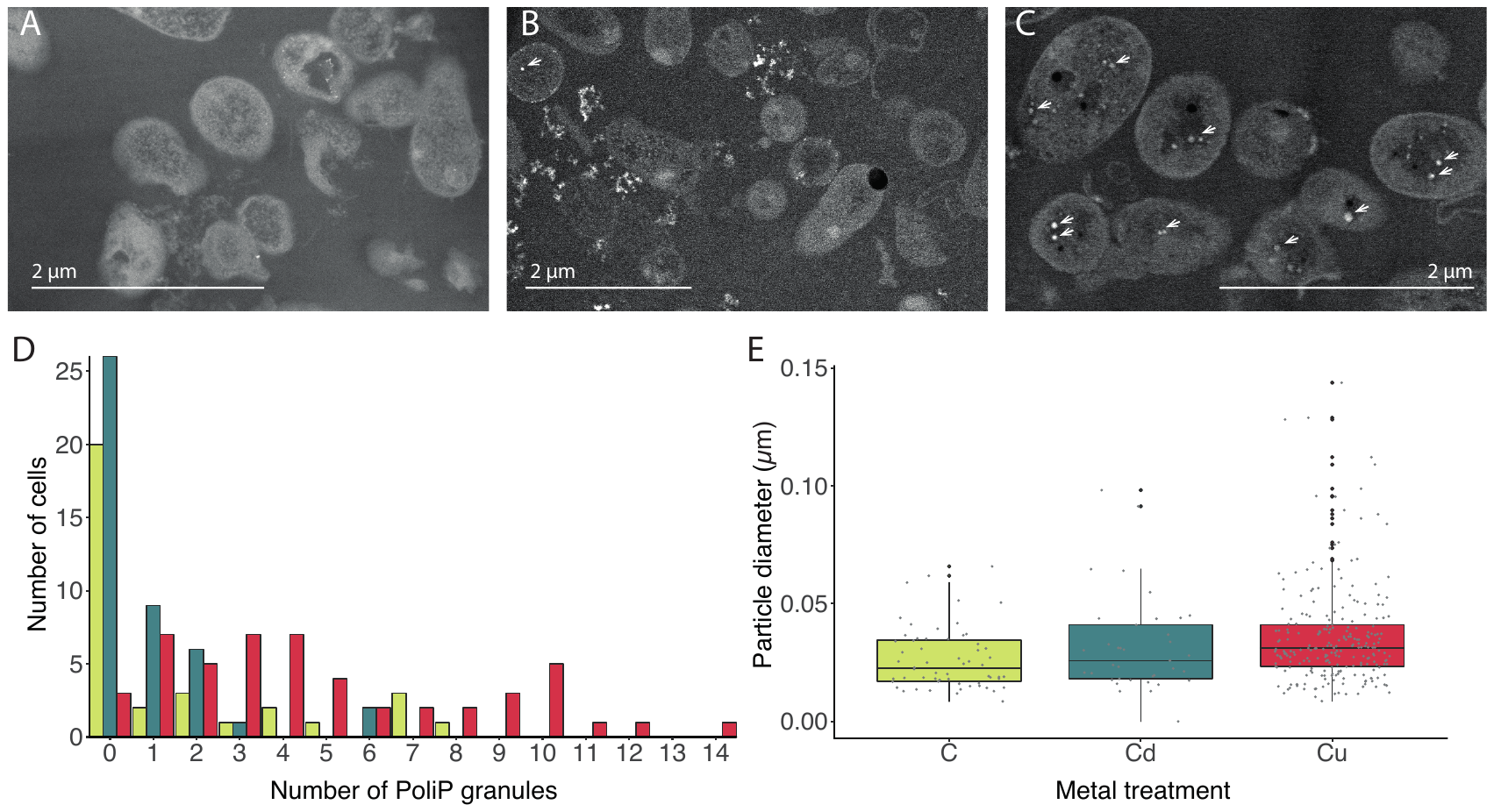
Figure 3
